# Expression of RcrB confers resistance to hypochlorous acid in uropathogenic *Escherichia coli*

**DOI:** 10.1101/2023.06.01.543251

**Authors:** Mary E. Crompton, Luca F. Gaessler, Patrick O. Tawiah, Lisa Pfirsching, Sydney K. Camfield, Colton Johnson, Kennadi Meurer, Mehdi Bennis, Brendan Roseberry, Sadia Sultana, Jan-Ulrik Dahl

## Abstract

To eradicate bacterial pathogens, neutrophils are recruited to the sites of infection, where they engulf and kill microbes through the production of reactive oxygen and chlorine species (ROS/RCS). The most prominent RCS is antimicrobial oxidant hypochlorous acid (HOCl), which rapidly reacts with various amino acids side chains, including those containing sulfur and primary/tertiary amines, causing significant macromolecular damage. Pathogens like uropathogenic *Escherichia coli* (UPEC), the primary causative agent of urinary tract infections (UTIs), have developed sophisticated defense systems to protect themselves from HOCl. We recently identified the RcrR regulon as a novel HOCl defense strategy in UPEC. The regulon is controlled by the HOCl-sensing transcriptional repressor RcrR, which is oxidatively inactivated by HOCl resulting in the expression of its target genes, including *rcrB*. *rcrB* encodes the putative membrane protein RcrB, deletion of which substantially increases UPEC’s susceptibility to HOCl. However, many questions regarding RcrB’s role remain open including whether *(i)* the protein’s mode of action requires additional help, *(ii) rcrARB* expression is induced by physiologically relevant oxidants other than HOCl, and *(iii)* expression of this defense system is limited to specific media and/or cultivation conditions. Here, we provide evidence that RcrB expression is sufficient to *E. coli*’s protection from HOCl and induced by and protects from several RCS but not from ROS. RcrB plays a protective role for RCS-stressed planktonic cells under various growth and cultivation conditions but appears to be irrelevant for UPEC’s biofilm formation.

**IMPORTANCE:** Bacterial infections pose an increasing threat to human health exacerbating the demand for alternative treatment options. UPEC, the most common etiological agent of urinary tract infections (UTIs), are confronted by neutrophilic attacks in the bladder, and must therefore be well equipped with powerful defense systems to fend off the toxic effects of RCS. How UPEC deal with the negative consequences of the oxidative burst in the neutrophil phagosome remains unclear. Our study sheds light on the requirements for the expression and protective effects of RcrB, which we recently identified as UPEC’s most potent defense system towards HOCl-stress and phagocytosis. Thus, this novel HOCl-stress defense system could potentially serve as an attractive drug target to increase the body’s own capacity to fight UTIs.

## INTRODUCTION

The human host uses oxidative stress as a strategy to limit bacterial infections (1, 2). For instance, during phagocytosis, neutrophils and macrophages produce high levels of reactive oxygen species (ROS) and reactive chlorine species (RCS) to kill ingested microbes (3, 4). To initiate the powerful oxidative burst, activated neutrophils release antimicrobial enzymes from granules into the phagosome, such as NADPH oxidase2 (NOX2) and myeloperoxidase (MPO) (5). The process starts with the assembly of NOX2 into the phagosomal membrane, where the enzyme reduces molecular oxygen into superoxide by oxidizing NADPH. Superoxide is then dismutated into hydrogen peroxide (H_2_O_2_) (6), a ROS that is fairly well tolerated by bacteria. However, MPO uses H_2_O_2_ as a substrate to catalyze the oxidation of halides (i.e. Cl^-^ and Br^-^) and pseudohalides (i.e. SCN^-^) into highly antimicrobial hypohalous acids (HOX). These include hypochlorous acid (HOCl), hypobromous acid (HOBr), and hypothiocyanous acid (HOSCN), respectively (5, 7, 8). The outcome of HOX production is determined by the differences in availability of the three anions as well as the distinct selectivity of MPO for each (pseudo-)halide. ∼70-80% of all H_2_O_2_ consumed by MPO yields HOCl and the remainder is accounted for by HOSCN, as the formation of HOBr is negligibly small due to the low bromide concentration available (9).

HOCl is characterized by its high oxidizing capacity with virtually any cellular macromolecule, including lipids, nucleic acid, and proteins (10–12). These oxidative modifications can cause protein aggregation, DNA strand cleavage, mis-metalation, ATP depletion, and a substantial reduction in free thiols, ultimately causing significant macromolecular damage and cell death. A common target of all HOX is the amino acid cysteine, which is subject to reversible (i.e. sulfenic acids; disulfide bonds) or irreversible thiol modifications (i.e. sulfinic and sulfonic acid) (3, 13–15). While irreversible thiol modifications can lead to protein aggregation and degradation, reversible thiol modifications often come along with significant structural and functional consequences that may play important roles for the cellular redox homeostasis (14–16). Notably, many bacteria utilize reversible cysteine modifications as a tool to activate appropriate response and defense systems against oxidative stress (15, 17). Moreover, HOCl reacts with primary and secondary amines (i.e. in arginine and lysine) resulting in the production of chloramines, which are also considered antimicrobial despite their four to five magnitudes lower reactivity compared to HOCl (18). Regardless, formation of these additional RCS further extends the oxidizing capabilities of HOCl on cellular macromolecules and illustrates the difficulty of dissecting their individual contributions to antimicrobial killing *in vivo* (12).

To mitigate the toxic effects of HOX, bacteria have evolved defense systems on both transcriptional and post-translational levels. Redox-sensing transcriptional regulators use conserved cysteine and/or methionine residues to modulate their activity (15, 17, 19, 20). Their HOX-mediated (in-)activation results in elevated transcript level of their target genes, many of which have been shown to protect the organism from ROS/RCS stress (18). Four HOCl-responsive transcriptional regulators have been identified in *Escherichia coli*. NemR is a transcriptional repressor that is inactivated by HOCl, resulting in the expression of *nemA*, an N-ethylmaleimide reductase, and *gloA*, a glyoxylase, both of which have been shown to contribute to HOCl resistance *in vitro* (21, 22). The HOCl-sensing transcriptional activator RclR induces the transcription of *rclABC* (23). While the function of RclB and RclC are still unknown, recent mechanistic studies shed light into RclA’s protective role for HOCl- and HOSCN-stressed *E. coli* (24, 25). While NemR and RclR sense HOCl through conserved cysteine residues, the transcriptional regulator HypT is activated through methionine oxidation and induces the transcription of genes involved in methionine and cysteine biosynthesis (26, 27). We recently reported that in comparison to *E. coli* K12 and enteropathogenic *E. coli* (EPEC) strains, uropathogenic *E. coli* (UPEC) are substantially more resistant to HOCl exposure and killing by neutrophils due to the activation of the RcrR regulon, an additional HOCl-defense system that many of the intestinal *E. coli* pathotypes lack (28). The UPEC RcrR regulon is regulated by the TetR-family transcriptional repressor RcrR, which becomes inactivated through reversible cysteine oxidation by HOCl resulting in the de-repression of the three structural genes *rcrARB*. UPEC’s increased HOCl resistance appears to be mediated by RcrB, a putative inner membrane protein of unknown function, as *rcrB*-deficient UPEC strains were as susceptible to HOCl as the HOCl-sensitive intestinal *E. coli* strains tested (28). However, questions regarding RcrB’s role remain open including whether the protein requires additional help for its protective effects. Recombinant expression of RcrB in *E. coli* strains that lack the operon suggest that RcrB is exclusively responsible for the increased HOCl resistance. We further provide evidence that RcrB expression is induced by and protects from several RCS, but not from ROS stress. RcrB plays a highly protective role for RCS-stressed planktonic cells under various growth and cultivation conditions but appears to be irrelevant for UPEC biofilm formation. Lastly, expression studies suggest a major role long-term role for RcrB, which cannot be complemented by its homolog RclC.

## RESULTS

### Recombinant expression of RcrB significantly improves HOCl resistance in RcrB-deficient *E. coli* strains

We recently reported that UPEC strains, including CFT073, an isolate from the blood of a patient with acute pyelonephritis (29), tolerate higher levels of HOCl compared to other *E. coli* pathotypes (28). We identified the protective gene cluster responsible for this phenotype, which is predominantly present in UPEC as well as in various invasive *E. coli* strains and consists of the three uncharacterized genes *rcrA, rcrR*, and *rcrB*. While we identified RcrR as an HOCl-sensing transcriptional repressor, the function of *rcrA* and *rcrB* remained unknown (28). Deletion of *rcrB* completely abolished CFT073’s increased resistance to HOCl, suggesting that the putative membrane protein RcrB is responsible for UPEC’s elevated HOCl resistance. To determine whether RcrB alone is indeed sufficient for the bacterial protection against HOCl and to examine whether its protective role extends beyond CFT073, we compared the growth behavior of different *E. coli* strains in the presence and absence of RcrB during sublethal HOCl stress. To reduce the possibility that media components react with and potentially quench HOCl, we performed our established lag phase extension (LPE)-based growth assay in MOPS-glucose (MOPSg) minimal media. We tested the impact of the presence and absence of RcrB in different strain backgrounds including strains that naturally lack *rcrB* by expressing plasmid encoded RcrB in the presence of various HOCl concentrations. Empty vector (EV) strains served as controls. Transformation of a Δ*rcrB* strain in the CFT073 background resulted in full complementation of the UPEC’s increased HOCl resistance **(Supplementary FIG 1A)**. Likewise, the same plasmid restored the HOCl resistance of the probiotic strain *E. coli* Nissle 1917 (EcN), a close relative of UPEC strain CFT073 (30), with an in-frame deletion in the gene homologous to *rcrB* (i.e. Δ*rcrB-H*) **(Supplementary FIG 1B)**. Taken together, these data suggest that leaky RcrB expression from the pET28a plasmid was sufficient to restore full HOCl resistance.

In our previous study, we showed that K-12 strain MG1655, which does not contain *rcrB,* is significantly more sensitive to HOCl than UPEC strain CFT073 (28). This is significant given that MG1655 is considered much more HOCl-resistant than most other non-pathogenic laboratory *E. coli* strains. We transformed the *rcrB*-encoding pET28a plasmid into MG1655 and performed our LPE assay in the presence and absence of increasing HOCl concentrations. As expected, MG1655 transformed with the empty vector control pET28a was substantially more sensitive to HOCl than the pET28a-containing UPEC strain CFT073 **(FIG 1A)**. However, leaky expression of pET28a-encoded RcrB resulted in LPE that were almost identical to CFT073 suggesting that RcrB alone and in the amount produced is sufficient to render MG1655 as resistant to HOCl as CFT073 **(FIG 1A)**. Like MG1655, the cystitis isolate UTI89, another UPEC model strain (31), also lacks *rcrB*, and its LPE under sublethal HOCl stress resembled that of MG1655 **(FIG 1B)**. Expression of pET28a-encoded RcrB in UTI89 resulted in substantially reduced LPE, which were even shorter than what we observed in pET28a-containing CFT073 **(FIG 1B)**. All in all, our experiments confirm the prominent role that RcrB has for protecting various *E. coli* strains from the deleterious effects of HOCl and indicate that the protein likely does so without any additional help.

**FIG 1.**
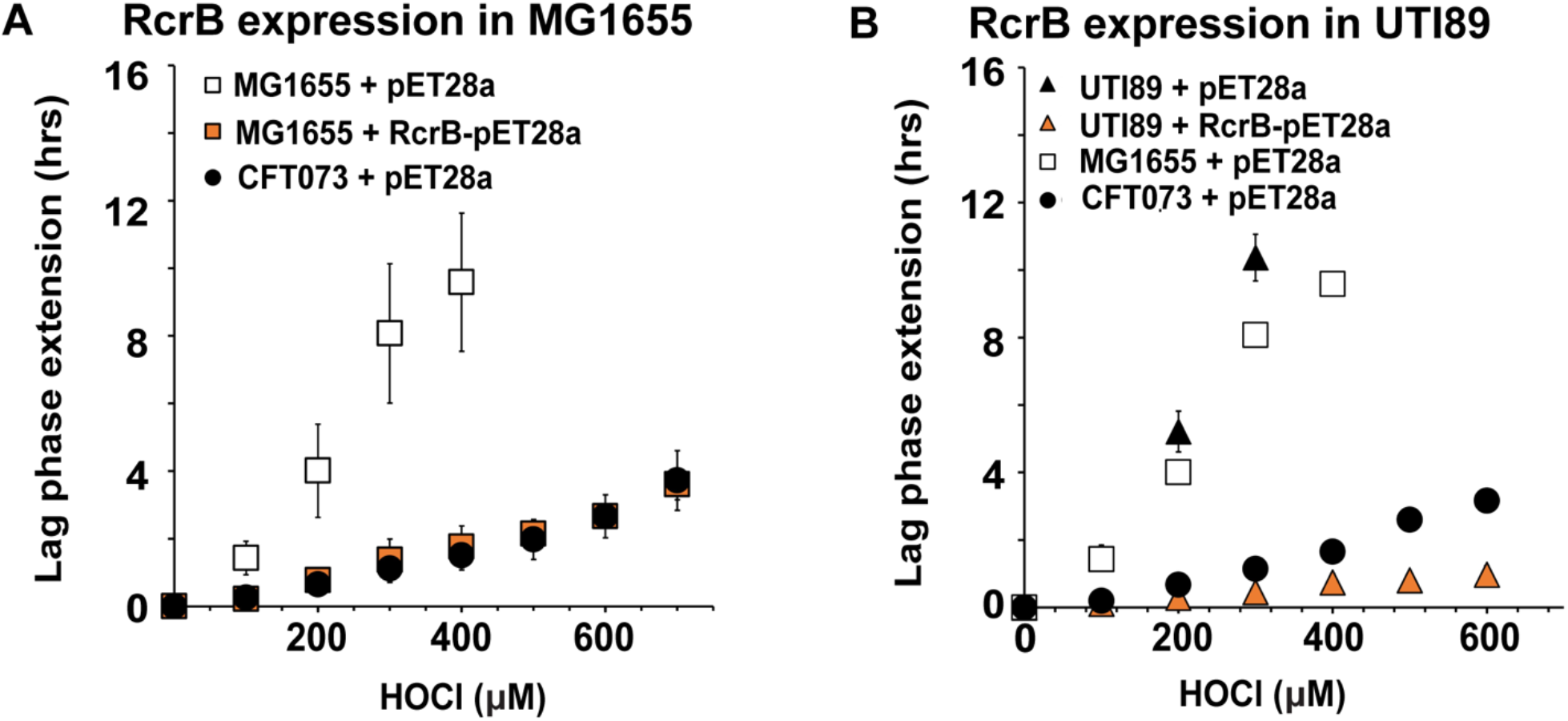
Recombinant expression of RcrB significantly improves HOCl resistance in different *E. coli* strains. Growth phenotype analyses of the K-12 *E. coli* strain MG1655 and UPEC strains CFT073 and UTI89 carrying pET28a or *rcrB*-pET28a were performed in MOPSg media in the presence of the indicated HOCl concentrations. HOCl-mediated LPE was calculated for each strain (see Materials and Methods for a detailed protocol). Squares, MG1655; circles, CFT073; triangles, UTI89; orange filling, + RcrB. Expression of RcrB in **(A)** MG1655 and **(B)** UTI89 resulted in an increased HOCl resistance and is comparable to what we observed in the CFT073 strain, *(n = 3-4, ± S.D.)*.

### The HOCl resistance of UPEC clinical isolates depends on the presence of RcrB

While both UTI89 and CFT073 were originally isolated from cystitis and pyelonephritis patients, respectively, they have been in the lab environment for decades and therefore may no longer serve as reliable representatives of UPEC infections. To further investigate the impact of RcrB on UPEC’s defense to HOCl stress, we used whole genome sequenced clinical UPEC isolates from asymptomatic patients, as well as those suffering from cystitis and pyelonephritis, respectively (32). We used quantitative real-time PCR (qRT-PCR) as a first test to determine whether these UPEC clinical isolates upregulate *rcrA* and *rcrB* upon exposure to sublethal HOCl stress. Both genes are members of the RcrR regulon, the expression of which was induced in HOCl-stressed CFT073 (28). We confirmed the transcriptional upregulation of both genes in VUTI229, VUTI288, and VUTI313, which are representative clinical isolates from pyelonephritis, cystitis and asymptomatic patients, respectively **(FIG 2A**). Next, we compared the growth of 13 sequenced clinical UPEC isolates at sublethal HOCl-stress to the HOCl-sensitive K-12 strain MG1655 and the HOCl-resistant UPEC strain CFT073. Our LPE assays revealed a strong correlation between increased HOCl resistance and the presence of *rcrB,* independent of whether the strain causes cystitis, pyelonephritis, or an asymptomatic infection (**FIG 2B-D**). The HOCl resistance profiles of strains carrying the *rcrB* gene in their chromosome (i.e. VUTI150/188/229/288/308/313/317/319/332) were similar to those observed in CFT073 (**FIG 2B-D**, *compare blue and black circles*). Similarly, strains that lack *rcrB* (i.e. VUTI207/334/369/409) resemble HOCl-sensitive *E. coli* strains such as MG1655 (**FIG 2B-D**, *compare red diamonds and white squares*).

**FIG 2.**
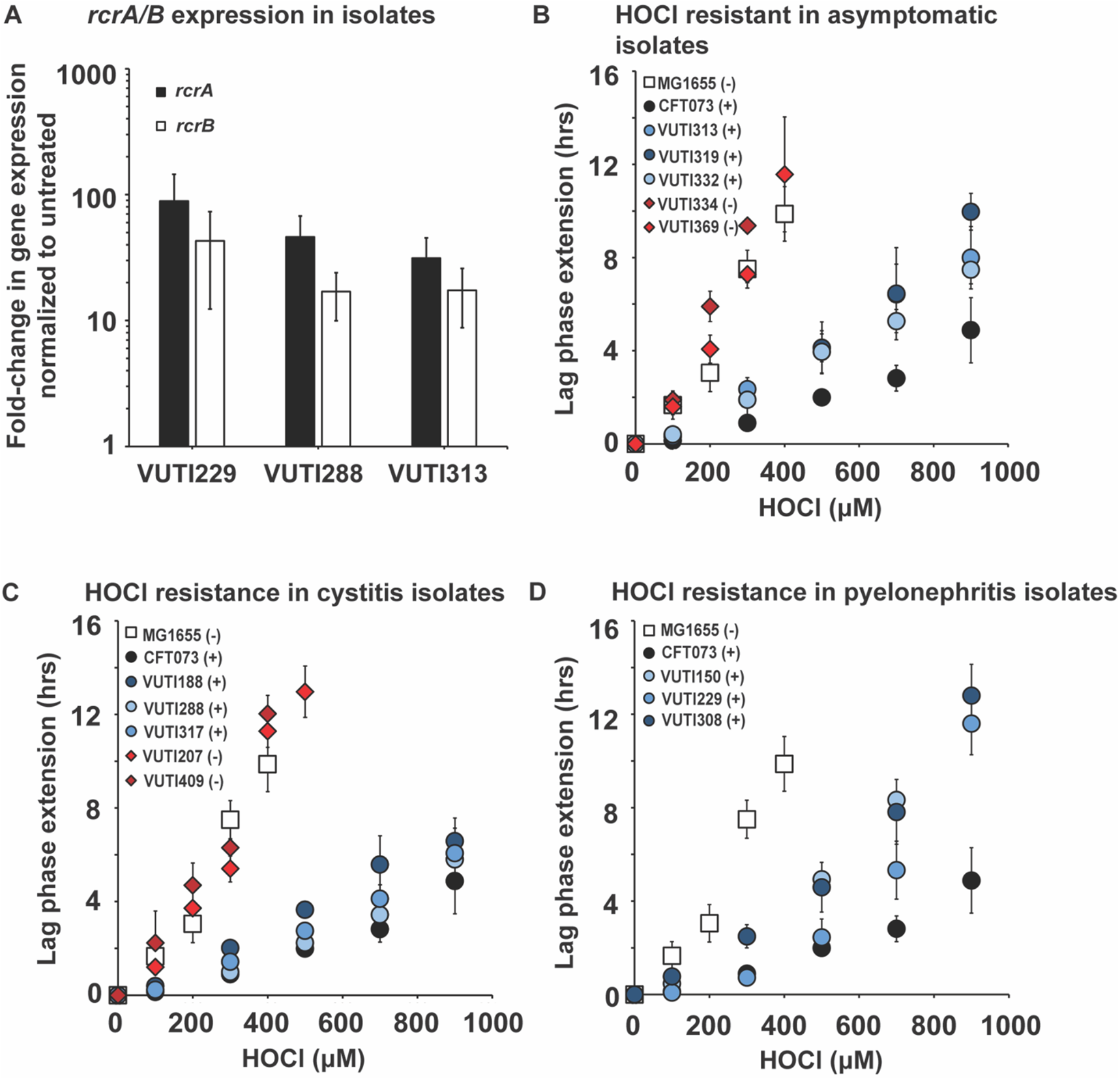
The HOCl resistance of UPEC clinical isolates depends on the presence of RcrB. **(A)** UPEC clinical isolates VUTI229, VUTI288, and VUTI 313 were grown to mid-log phase in MOPSg and changes in the expression of *rcrA* (black bars) and *rcrB* (white bars) upon 15 min exposure to 1 mM HOCl were determined by qRT-PCR. *rcrA* and *rcrB* mRNA level were elevated in all three strains in the presence of HOCl *(n = 3, ± S.D.)*. **(B-D)** Growth phenotype analyses of the indicated UPEC clinical isolates causing **(B)** an asymptomatic infection, **(C)** cystitis, and **(D)** pyelonephritis were performed in MOPSg media in the presence of the indicated HOCl concentrations and compared to the K-12 *E. coli* strain MG1655 and UPEC strain CFT073. HOCl-mediated LPE was calculated for each strain (see Materials and Methods for a detailed protocol). The presence (+) and absence (-) of the *rcrB* gene in the genome of each strain is indicated in the legend. Diamonds, clinical isolates lacking *rcrB*; circles, UPEC carrying *rcrB*; squares, MG1655. The increased HOCl resistance of UPEC clinical isolates depends on the presence of *rcrB*, *(n= 4-7, ± S.D.)*.

### RcrB plays an important role for UPEC’s HOCl resistance under various growth conditions

In nature, bacteria often experience nutrient-limiting conditions, and their growth more likely resembles stationary phase cells in batch culture. Stationary phase and non-growing, dormant bacteria are typically more resistant to stress compared to exponentially growing cells, which is attributed to differences in their gene regulation and metabolic activities (33–35). Thus, we were wondering whether the cellular impact of RcrB during HOCl stress differs under various growth conditions. We performed LPE-based growth analyses of exponentially growing CFT073 and Δ*rcrB* cells in presence of increasing HOCl concentrations and compared the outcome to cells that were in the late logarithmic (log)/early stationary phase of growth. We found that independent of the growth phase, Δ*rcrB* cells exhibited significantly higher sensitivity to HOCl as evidenced by their longer LPE in comparison to the wildtype (**FIG 3A, B**). We made very similar observations in experiments with different cell densities upon which HOCl was added (**FIG S2**). Interestingly, we observed a declining difference in HOCl sensitivity between the wild-type and mutant strains at higher starting optical densities (**FIG S2**). We further performed survival analyses for both the exponential and stationary cultures of each strain under growing and non-growing conditions to test whether metabolically active and inactive bacteria rely on RcrB differently. We cultivated both strains in MOPSg, washed the cells, and shifted one part into fresh MOPSg media (containing glucose) to allow for continuous growth prior to the exposure to HOCl. We transferred the other part into fresh MOPS media without glucose (MOPS), which resulted in an immediate growth arrest. We reasoned that this approach should allow us to keep the media composition constant while only varying the growth status of the bacteria. Under all the conditions tested, cells that lack RcrB (i.e. Δ*rcrB*) showed between 2- and 5-log higher sensitivity to HOCl (**FIG 3C, D**). The least difference in survival between the wildtype and Δ*rcrB* mutant was observed upon shift of late log/stationary cells to MOPS resulting in only 1-log difference in survival (**FIG 3D**). In summary, our data indicate that RcrB is an important player in UPEC’s HOCl response independent of the bacterial growth status.

**FIG 3.**
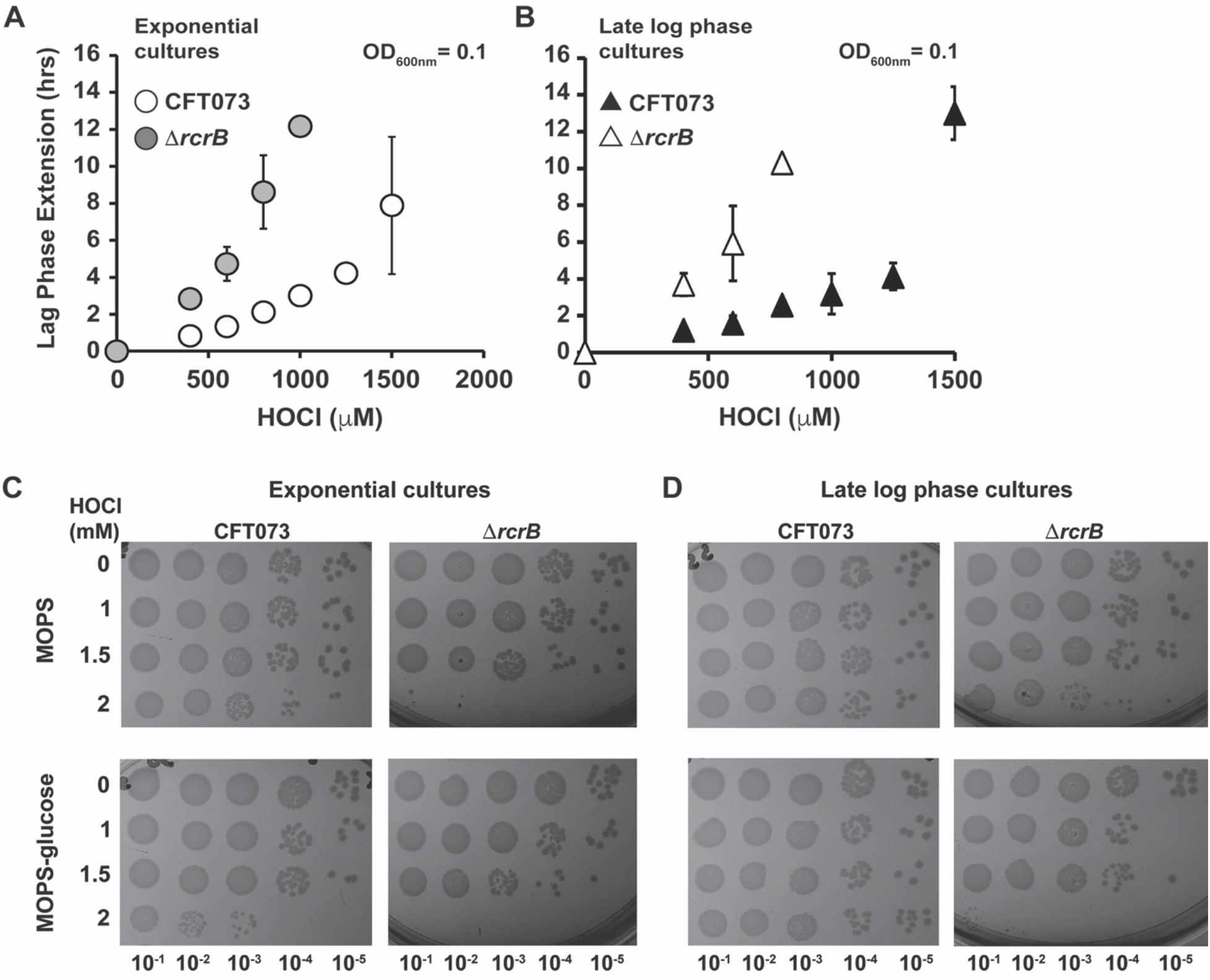
RcrB plays an important role for UPEC’s HOCl resistance under various growth conditions. **(A, B)** Growth phenotype analyses of UPEC strains CFT073 and Δ*rcrB* were performed in MOPSg media in the presence of the indicated HOCl concentrations. HOCl-mediated LPE was calculated for each strain (see Materials and Methods for a detailed protocol). **(A)** *Exponential cultures (left):* overnight cultures were diluted into fresh MOPSg to an OD_600_ = 0.01 and grown until they reached an OD_600_ = 0.1 before they were split up and cultivated in the presence of the indicated HOCl concentrations. **(B)** *Late log cultures (right):* overnight cultures were diluted 25-fold into fresh MOPSg and grown until they reached late log / early stationary phase (OD_600_ ∼2) before they were diluted again to OD_600_ = 0.1 and cultivated in the presence of the indicated HOCl concentrations; *(n = 3-4, ± S.D.)*. **(C, D)** Effects of HOCl on the survival of growing and non-growing UPEC strains CFT073 and Δ*rcrB.* CFT073 and Δ*rcrB* were grown in MOPSg media to **(C)** early exponential phase (OD_600_ = 0.1) or **(D)** late log / early stationary phase (OD_600_ ∼2) before they were shifted to MOPS media in the presence or absence of glucose. The indicated concentrations of HOCl were added. After 145 min of incubation, excess HOCl was quenched by the addition of 5-fold excess thiosulfate, cells were serially diluted with PBS, spot-titered onto LB agar plates, and incubated overnight at 37°C. The experiments were repeated at least three independent times.

### RcrB does not protect from HOCl stress when UPEC is grown as biofilms

The formation of biofilms is a robust response of bacteria to switch from a planktonic and motile to a sessile lifestyle, which provides UPEC isolates with increased bladder fitness and antimicrobial resistance (36). UPEC biofilms are also formed on abiotic surfaces such as catheters, increasing the risk for the development of catheter associated UTIs. Recent studies provided evidence for an increased tendency of bacteria to switch from planktonic to the biofilm lifestyle when they encounter sublethal HOCl stress, likely as an adaptive stress response (15, 37–39). To further investigate RcrB’s role for UPEC’s increased HOCl resistance, we compared biofilm formation of UPEC wildtype and Δ*rcrB* cells in presence of HOCl. The two strains were grown in LB without salt for 6 hours to facilitate initial attachment to the well before the incubation was resumed for another 16 hours in the presence of different concentrations of HOCl. We quantified the biofilm formed on the liquid-solid interface using crystal violet (CV) staining. While we found ∼30% increased biofilm formation upon exposure of both strains to 4 mM HOCl, neither the presence nor absence of HOCl resulted in significant differences in CV staining between both strains (**FIG 4**), suggesting that RcrB has no influence on HOCl-mediated biofilm responses or biofilm formation in general.

**FIG 4.**
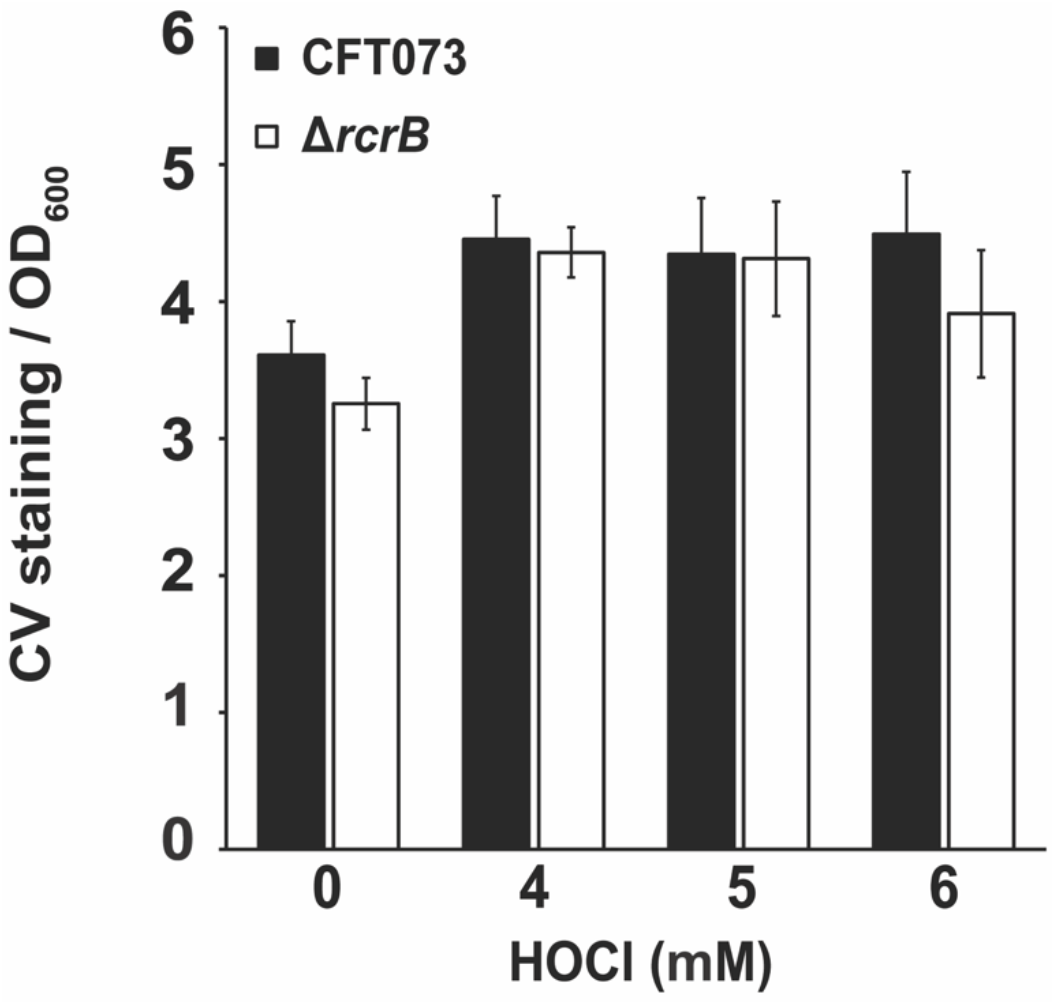
RcrB does not protect from HOCl stress when UPEC is grown as biofilms. Overnight cultures of CFT073 (black bars) and Δ*rcrB* (white bars) were diluted into LB without NaCl and cultivated under static conditions in 96-well plates in the absence and presence of the indicated HOCl concentrations. After 16 hours, OD_600_ was determined, and planktonic cells removed. The biofilms were washed twice with PBS, stained with 0.1% crystal violet (CV) for 15 min, washed again twice with PBS to remove unbound CV dye. CV bound to biofilm cells was extracted with 30% (v/v) glacial acetic acid and OD_595_ was determined. CV staining (OD_595_) was normalized to the OD_600_ of planktonic cells, *(n = 2 [with 8 technical replicates each], ± S.D.)*.

### RcrB protects UPEC from HOCl stress when cultivated in artificial urine media (AUM)

We have shown that RcrB provides protection from HOCl-stress when the cells were cultivated in MOPSg, a minimal media used to reduce the possibility that media components react with and potentially quench HOCl. However, this environment may not necessarily resemble the physiological conditions in the urinary tract although UPEC is considered extremely versatile with regards to the different environments it can inhabit. UPEC strains grow as harmless commensals in the nutrient- and carbon-rich human gastrointestinal tract but transition rapidly into a pathogenic lifestyle upon entry into the urinary tract, an environment that is characterized by its high osmolarity and nitrogen abundance but also lower nutrient content and bacterial competition (40). Thus, we sought to confirm our findings in artificial urine media (AUM), which resembles more closely the conditions UPEC typically encounter in the urinary tract (41). MG1655, CFT073, and Δ*rcrB* grown in AUM were treated with 380 μM HOCl, a concentration that resulted in a LPE of ∼ 6 hours for the CFT073 wildtype **(FIG 5A)**. Consistent with data in MOPSg, LPE were significantly increased by HOCl in the Δ*rcrB* and MG1655 strains, indicating that the presence of RcrB in CFT073 wildtype contributes to a better recovery from the HOCl exposure **(FIG 5A)**. To verify that an extended lag phase in HOCl-treated MG1655 and Δ*rcrB* cells correlates with decreased survival, we performed time killing assays, in which we exposed the three strains to 668 μM HOCl, a concentration that resulted in ∼25% reduction in CFT073 wildtype survival (**FIG 5B**). The absence of *rcrB*, either in CFT073 (i.e. Δ*rcrB*) or in MG1655, caused a much more drastic effect and resulted in a ∼75% survival decline **(FIG 5B)**. To confirm an analogous effect in clinical UPEC isolates, we performed overnight growth assays in AUM focusing on strains that either contain *rcrB* (i.e. VUTI308) or lack the gene (i.e. VUTI207). Cultures of each clinical isolate were allowed to grow in a 96-well plate overnight in the presence and absence of increasing concentrations of HOCl, and the optical density at 600 nm was recorded after 18 hours. Upon exposure to 350 µM HOCl, growth of the RcrB-containing strain VUTI308 was unaffected, whereas VUTI207 showed a drastic reduction in optical density **(FIG 5C)**. Therefore, we conclude that RcrB’s protective role from HOCl stress is also prominent in AUM.

**FIG 5.**
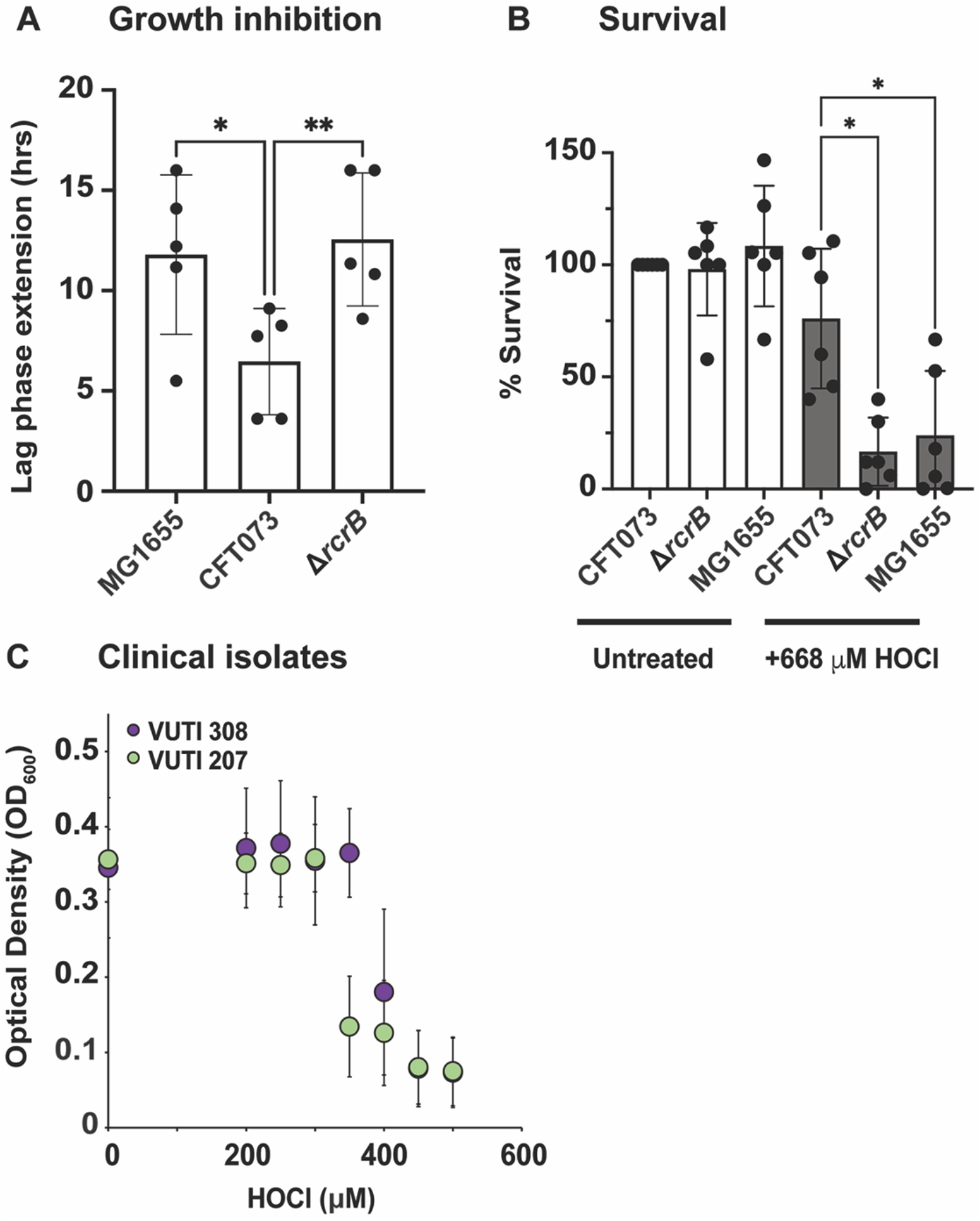
RcrB protects UPEC from HOCl stress when cultivated in artificial urine media (AUM) **(A)** Growth phenotype analyses of the K-12 *E. coli* strain MG1655 and UPEC strains CFT073 and Δ*rcrB* were performed in AUM media in the presence of 380 μM HOCl. HOCl-mediated LPE was calculated for each strain (see Materials and Methods for a detailed protocol). HOCl exposure in MG1655 and Δ*rcrB* caused ∼6 hours higher LPE compared to CFT073, *(n = 5, ± S.D.).* **(B)** Survival of MG1655, CFT073, and Δ*rcrB* upon HOCl exposure was examined in a time-killing assay. Overnight strains grown in LB were diluted ∼25-fold into AUM (OD_600_ = 0.1) and exposed to 668 μM HOCl. After 60 min of incubation, cells were serially diluted in PBS, spot-titered onto LB agar, and incubated overnight for CFU counts. Survival of MG1655 and Δ*rcrB* were significantly reduced compared to CFT073, *(n= 6, ± S.D.).* **(C)** UPEC clinical isolates VUTI308 (possesses chromosomal *rcrB*) and VUTI207 (chromosomal *rcrB* absent) were grown in AUM in the presence of the indicated HOCl concentrations. After 16 hours of growth at 37°C, OD_600_ was measured and plotted against the HOCl concentrations tested. VUTI308 showed growth between 300-400 μM HOCl exposure while VUTI207 did not, *(n= 4, ± S.D.)*.

### Transcription of the RcrR regulon is induced over a longer time following exposure to HOCl

Our previous RNAseq transcriptomic analyses identified the members of the RcrR regulon among the most upregulated genes in HOCl-stressed CFT073 cells (28). To better understand the factors that result in the induction of the RcrR regulon, we aimed to *(i)* examine the range of HOCl concentrations that result in transcriptional induction of *rcrARB*, *(ii)* determine the kinetics of *rcrARB* upregulation, and *(iii)* compare their expression to members of the NemR and RclR regulons, two additional HOCl-defense systems that contribute to increased survival in HOCl-stressed *E. coli* (21, 23, 25). We cultivated CFT073 in MOPSg to mid-log phase and exposed the cells to increasing concentrations of HOCl for 15 min. We used qRT-PCR to analyze the transcript levels of each member of the RcrR regulon. The addition of HOCl concentrations between 0.1 and 2.5 mM resulted in similar mRNA level for all three genes, which were between 20- and 75-fold higher than in untreated CFT073 cells suggesting that very little HOCl is already sufficient to mount a response (**FIG 6A**). Treatment with 5 mM HOCl did not result in a significant upregulation of gene expression for any of the three genes, likely because the concentration was no longer sublethal and caused substantial killing **(Supplementary FIG S3A**). A time course analysis upon exposure of CFT073 to 1 mM HOCl, a sublethal concentration that does not cause killing, revealed that *rcrARB* induction remains elevated over at least 45 min (**FIG 6B, Supplementary FIG S3b**). Likewise, the expression of *nemR* and *nemA*, a member of the NemR regulon, as well as *rclR* and *rclA*, a gene under the control of the HOCl-sensing transcriptional activator RclR, were also rapidly induced by HOCl (**FIG 6C-D**). However, their transcript level declined rapidly over 45 min, which is consistent with previous studies in the non-pathogenic *E. coli* strain MG1655 (21, 42). These results further indicate that the presence of the RcrR regulon, particularly RcrB, is needed for the cellular protection for longer time periods in comparison to the other two defense systems.

**FIG 6.**
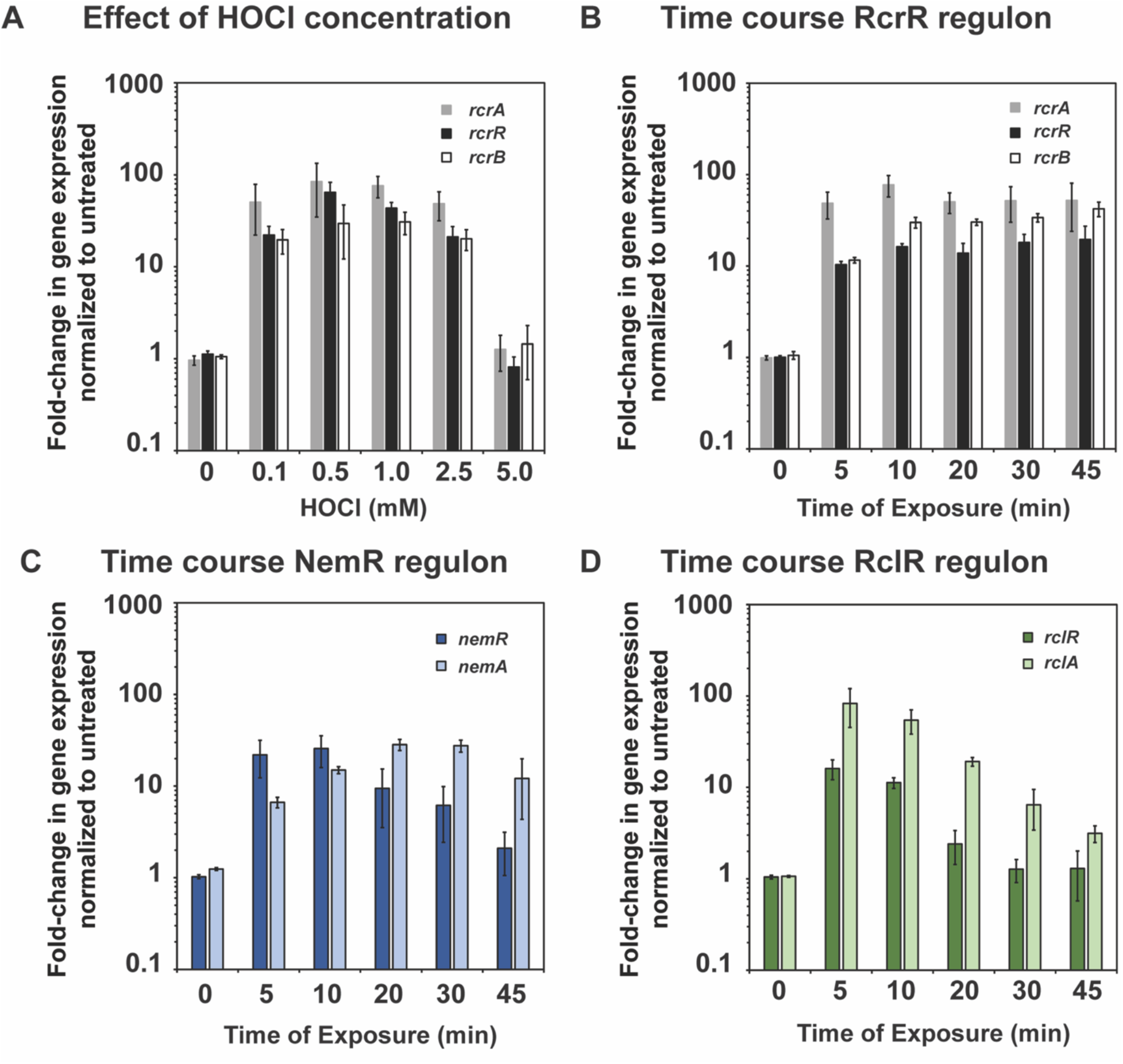
Transcription of the RcrR regulon is induced over a longer time following exposure to HOCl. UPEC strain CFT073 was grown in MOPSg to mid-log phase (OD_600_ ∼0.5-0.55) before it was incubated with the indicated HOCl concentrations for the indicated time. Transcription was stopped by the addition of ice-cold methanol. RNA was extracted, residual DNA removed, and mRNA reverse transcribed into cDNA. Transcript levels of the indicated genes were determined by qRT-PCR. Gene expression was normalized to the housekeeping gene *rrsD* and calculated as fold changes based on expression levels in the untreated control. **(A)** Fold-change in transcript level of select members of the RcrR regulon following treatment with increasing concentrations of HOCl for 15 min, (*n=3, ± S.D.*). **(B-D)** Fold-changes in transcript level of select members of the **(B)** RcrR, **(C)** NemR, and **(D)** RclR regulons following treatment with 1 mM HOCl for the indicated time points, (*n=3, ± S.D.*).

### Transcription of the RcrR regulon is induced in the presence of HOCl and its byproducts but not H_2_O_2_ or HOSCN

Next, we examined whether the induction of *rcrARB* transcription is specific to HOCl or can also be induced by any of the other ROS/RCS that are either produced in the neutrophil phagosome during oxidative burst (i.e. HOBr, HOSCN, H_2_O_2_) or are byproducts from the reaction of HOCl with different biological compounds (NCT, NH_2_Cl, Ca(OCl)_2_, Gly-Cl). Using qRT-PCR, we analyzed the fold-change in *rcrR*/*rcrB* gene expression following exposure to sublethal concentrations of each oxidant, which we had determined in survival assays before. Except for H_2_O_2_ and HOSCN, which resulted in insignificant transcriptional changes or a 5-fold down-regulation, respectively, exposure to all other stressors resulted in an upregulation of both *rcrR* and *rcrB* that was comparable to what we observed with HOCl **(FIG 7)**. These results indicate that expression of RcrR regulon is stimulated by RCS in general but not by ROS.

**FIG 7.**
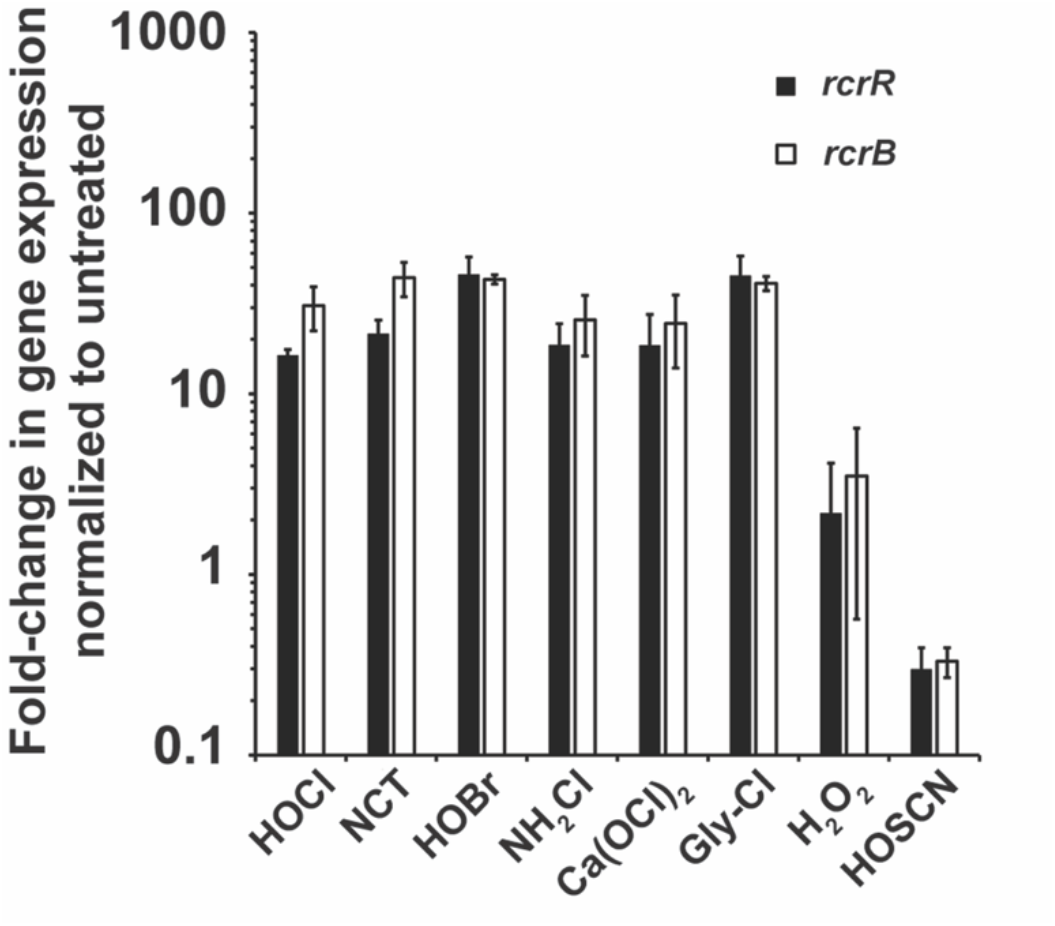
Transcription of the RcrR regulon is induced in the presence of HOCl and its byproducts but not H_2_O_2_ or HOSCN. UPEC strain CFT073 was grown in MOPSg to mid-log phase (OD_600_ ∼0.5-0.55) prior to incubation with sublethal concentrations of the indicated stressors for 15 min (HOCl, 1 mM; NCT, 0.3 mM; NH_2_Cl, 0.3 mM; HOSCN, 0.35 mM; HOBr, 0.5 mM; Ca(OCl)_2_, 0.2 mM; Gly-Cl, 0.5 mM; H_2_O_2_, 10 mM). Transcription was stopped by the addition of ice-cold methanol. RNA was extracted, residual DNA removed, and mRNA reverse transcribed into cDNA. The induction of transcript levels of the indicated genes was determined by qRT-PCR. Gene expression was normalized to the housekeeping gene *rrsD* and calculated as fold changes based on expression levels in the untreated control. *rcrB* and *rcrR* transcript level were increased following treatment with HOCl, NCT, HOBr, NH_2_Cl, Ca(OCl)_2_, and Gly-Cl, respectively. No change in transcript level was seen in the presence of H_2_O_2_, while exposure to HOSCN resulted in downregulation of *rcrB*/*rcrR* (*n=3, ± S.D.*).

### The RcrR regulon protects from HOCl/HOBr stress and its byproducts but not from H_2_O_2_ and HOSCN

Given that the expression of the RcrR regulon is induced by several RCS, we next examined whether these findings correlate with improved survival. Using our growth curve-based assays, we exposed CFT073, Δ*rcrB* (lacks RcrB), and Δ*rcrR* (constitutively expresses RcrB) to increasing concentrations of the indicated oxidants and recorded their growth. Likewise, we followed the same set-up using MG1655 (lacks RcrB), which was transformed with the RcrB-expressing plasmid (i.e. MG1655 + *rcrB*-pET28a), and compared its growth to MG1655 cells containing the empty vector control (i.e. MG1655 + pET28a). Consistent with our qRT-PCR data **(FIG 7)**, we neither observed significant differences in growth and/or LPE between the three strains in the CFT073 background nor between those in the MG1655 backgrounds when cells were treated with HOSCN and H_2_O_2_, respectively **(FIG 8, Supplementary FIG S4-5).** In contrast, exposure of NCT and Gly-Cl revealed a higher susceptibility of strains that lack RcrB (i.e. Δ*rcrB* and MG1655+pET28a) compared to those that express the membrane protein (i.e. CFT073 wildtype and MG1655+*rcrB*-pET28a) **(FIG 8, Supplementary FIG S4-5)**. However, the differences in growth deficit between these strains were smaller than what we observed upon HOCl exposure. Constitutive expression of RcrB, as it occurs in the Δ*rcrR* strain, provides additional protection from HOCl, Gly-Cl, and NCT, respectively. In summary, our results indicate that the RCS-induced expression of RcrR regulon provides substantial protection from RCS in general but not from ROS.

**FIG 8.**
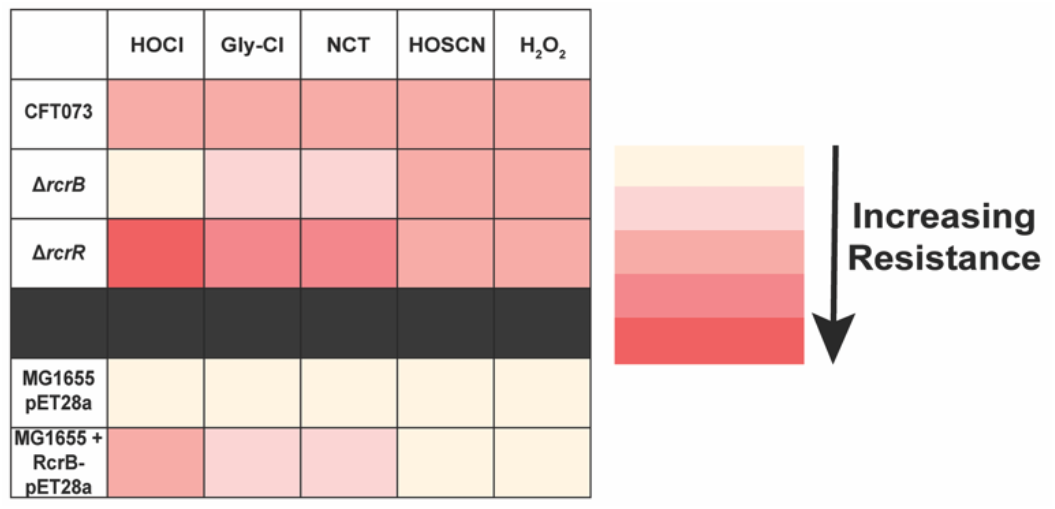
The RcrR regulon protects from HOCl stress and its byproducts but not from H_2_O_2_ and HOSCN. Growth phenotype analyses of the K-12 *E. coli* strain MG1655 carrying pET28a or *rcrB*-pET28a, respectively, as well as UPEC strains CFT073, Δ*rcrB*, and Δ*rcrR* were performed in MOPSg media in the presence of the indicated stressor concentrations. LPE were calculated for each strain (see Materials and Methods for a detailed protocol). Endogenous and recombinant expression of RcrB in CFT073 and MG1655, respectively, resulted in an increase in resistance to all stressors tested except to HOSCN and H_2_O_2_.

### RcrB and RclC do not complement each other during HOCl exposure

RcrB encodes a hypothetical protein, which according to our bioinformatic analysis shares partial homology with the membrane protein RclC, a member of the HOCl-sensing RclR operon with unknown function (23). Therefore, we examined whether the recombinant expression of RcrB and RclC can complement each other in CFT073 strains defective in either *rcrB* (i.e. Δ*rcrB*), *rclC* (i.e. Δ*rclC*) or both genes (i.e. Δ*rcrB*Δ*rclC*) during HOCl stress **(FIG 9)**. Empty vector (EV) strains served as controls. We compared the growth behavior of the different strains after exposing them to sublethal HOCl concentrations. In contrast to Δ*rcrB,* which as expected showed substantially higher LPE than the wildtype cells (**FIG 9**, *compare black and grey circles*), no such difference was observed for Δ*rclC* (**FIG 9**, *compare black circles and white diamonds*). Moreover, LPE were not significantly increased in Δ*rcrB*Δ*rclC* compared to Δ*rcrB* (**FIG 9**, *compare white and grey circles*), suggesting that in contrast to MG1655, RclC may not play an important role for HOCl resistance in UPEC. Consistent with these data, recombinant expression of RclC had no positive effect on the growth of Δ*rcrB* (**FIG 9**, *compare grey and grey-striped circles*), while expression of RcrB in the Δ*rclC* mutant resulted in substantial resistance to HOCl, as evidence by no to very little LPE of this strain (**FIG 9**, *compare white and grey diamonds*).

**FIG 9.**
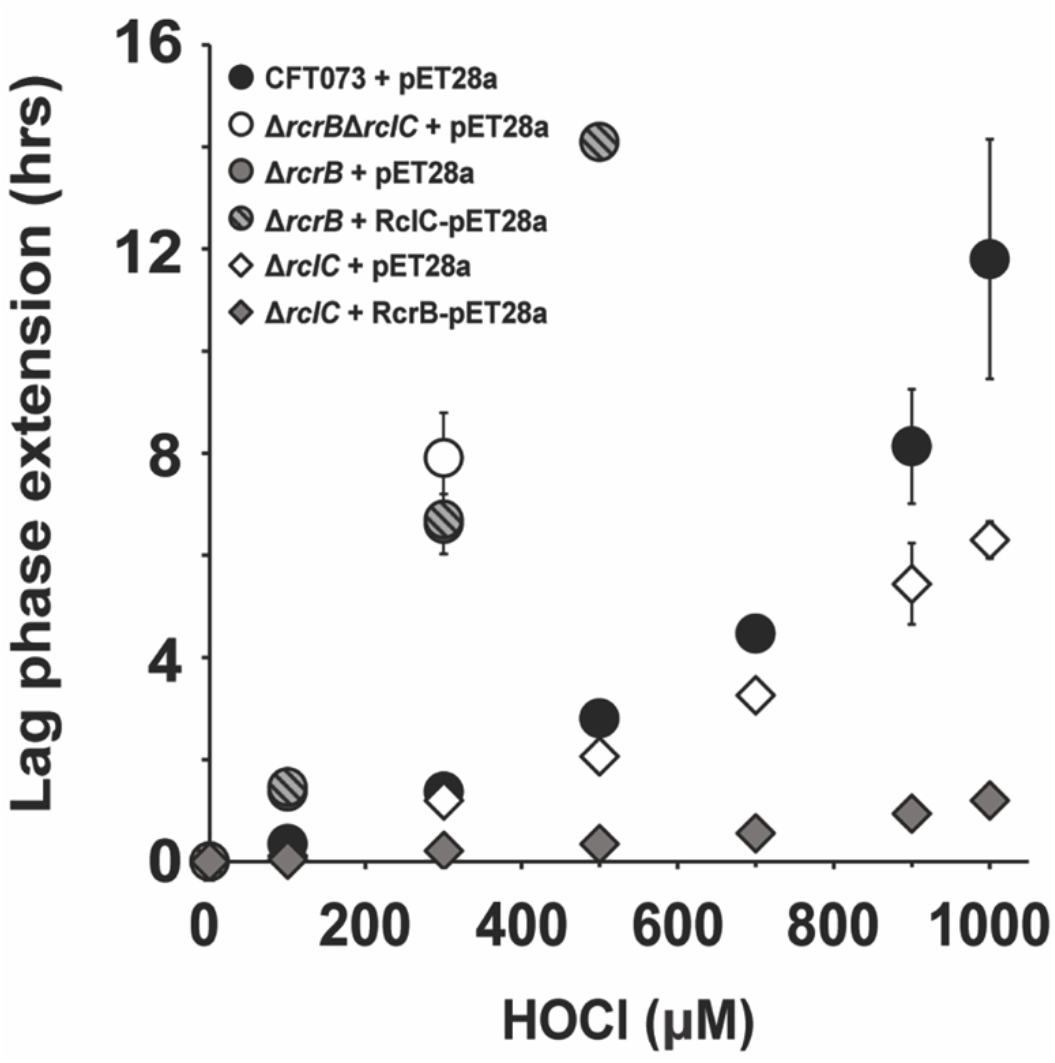
RcrB and RclC do not complement each other during HOCl exposure. Complementation analyses of the UPEC strains CFT073, Δ*rcrB*, Δ*rclC*, and Δ*rcrB*Δ*rclC* in the presence of *rcrB*-pET28a, *rclC*-pET28a, and the empty vector control pET28a were performed in MOPSg media in the presence of the indicated HOCl concentrations. HOCl-mediated LPE was calculated for each strain (see Materials and Methods for a detailed protocol). The absence of CFT073 *rclC* does not cause increased HOCl sensitivity nor does recombinant RclC expression complement the Δ*rcrB* strain, *(n = 3, ± S.D.)*.

## DISCUSSION

In this study, summarized in **FIG 10**, we characterized the significance of the RcrR regulon, comprised of the three genes *rcrA*, *rcrR*, and *rcrB,* for UPEC’s survival during RCS stress. We provide evidence that the elevated expression of *rcrARB* provides protection against HOCl, the major RCS, but also to HOBr and chlorinated byproducts that HOCl forms with taurine, glycine, ammonium, and calcium, respectively (43–46). In contrast, no *rcrARB* upregulation was observed in the presence of thiol-specific ROS (i.e., H_2_O_2_) and the hypohalous acid HOSCN. The protective effect of the RcrR regulon is exclusively mediated by RcrB given that expression of plasmid encoded RcrB is sufficient to significantly increase the HOCl resistance of *E. coli* strains that lack the RcrR regulon. RCS induces RcrB expression in both exponentially growing and stationary phase UPEC cells cultivated in minimal and artificial urine media, respectively. However, no such RcrB-mediated protective effect was observed when UPEC was present as a biofilm. In contrast to members of the NemR and RclR regulons, which provide protection against HOCl in *E. coli* K12 strains, expression of the RcrR regulon appears to be the main RCS defense system in UPEC given its prolonged expression during RCS exposure and the lack of complementation between RcrB and its homolog RclC (**FIG 10**).

**FIG 10.**
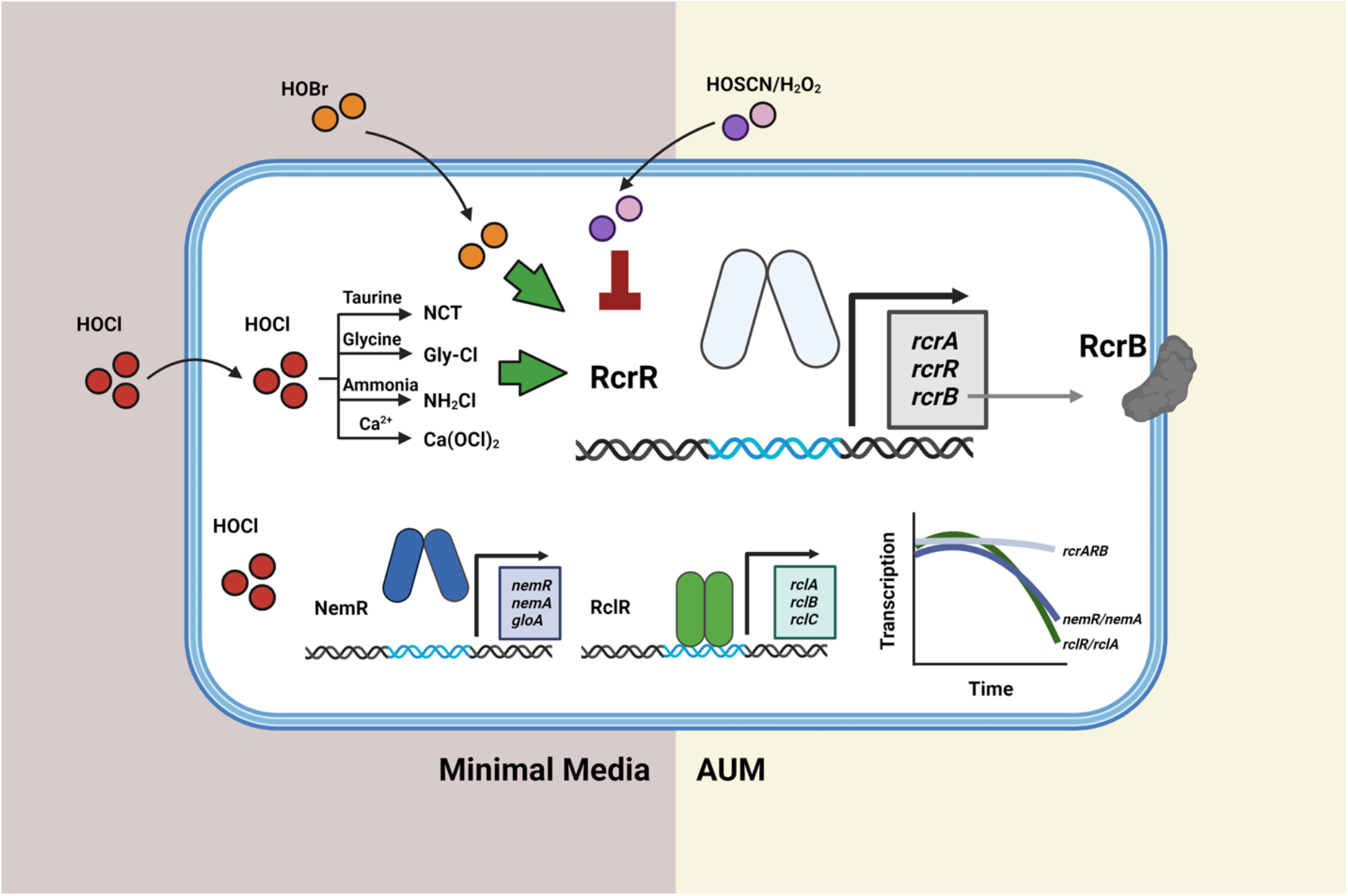
Summary of the findings of this study. The RcrR regulon plays an important role for UPEC’s survival during RCS stress, including their exposure to HOCl, HOBr, and the byproducts of HOCl, such as NCT, Gly-Cl, NH_2_Cl, and Ca(OCl)_2_, respectively. The protective effect of the RcrR regulon is exclusively mediated by RcrB given that expression of plasmid encoded RcrB is sufficient for an increased HOCl resistance in *E. coli* strains that lack the RcrR regulon. In contrast, RcrR neither responds to HOSCN nor to H_2_O_2_. RCS induces RcrB expression in exponentially growing and stationary phase UPEC in minimal and artificial urine media. No such protective effect was observed in biofilm-forming UPEC cells. In contrast to the NemR and RclR regulons which play significant roles for protection of *E. coli* K12 strains from HOCl, expression of the RcrR regulon appears to be the main RCS defense system in UPEC given its prolonged expression during RCS exposure.

*rcrARB* is controlled by the HOCl-sensing transcriptional repressor RcrR, which represses *rcrARB* transcription under non-stress conditions. However, HOCl negatively affects RcrR’s transcriptional repressor activity by reversibly oxidizing two conserved cysteine residues (28). As a result of conformational changes, RcrR dissociates from the promoter and *rcrARB* transcript level increase. RcrB appears to be the major driver for UPEC’s increased resistance to HOCl *in vitro* and phagocytosis by activated neutrophils (28). The *rcrARB* operon is primarily found in members of invasive *E. coli* pathotypes, including adherent-invasive *E. coli,* avian pathogenic *E. coli,* and UPEC (28). In contrast, non-invasive *E. coli*, such as the enteropathogenic strain O127:H6 and the K-12 strain MG1655, lack the presence of the gene cluster and were shown to be significantly more sensitive to HOCl. However, recombinant expression of RcrB rendered MG1655 significantly more resistant to HOCl, similar to what we observed for the RcrB-containing UPEC strain CFT073 (**FIG 1**). Likewise, the HOCl resistance of UPEC strain UTI89, a cystitis isolate that lacks the RcrR regulon (31), was significantly higher when the cells were transformed with the RcrB expression plasmid. When the *rcrB* homolog was deleted from the probiotic *E. coli* strain Nissle 1917, the sensitivity to HOCl became more profound, resembling what we observed in MG1655, *rcrB*-deficient CFT073 cells, or clinical isolates that lack the gene cluster (**FIG 1&2**). *E. coli* is an important member of the mammalian intestine and characterized by its great diversity. While most *E. coli* strains colonize the gut as harmless commensals, several *E. coli* pathotypes have evolved that cause a wide variety of intestinal and extraintestinal diseases (47). Among the group of extraintestinal *E. coli* are UPEC, the primary causative agent of UTIs (47, 48). UTIs are considered one of the most common bacterial infections worldwide and affect ∼150 million people, primarily women at all ages, each year (49). Different forms of UTIs exist depending on the location of the infection, such as cystitis, where UPEC infection occurs in the bladder, or pyelonephritis in the kidney (47). However, prior to successful colonization in the bladder, activated neutrophils infiltrate the bladder to clear bacterial intruders such as UPEC through a powerful oxidative burst (48). Oxidative burst occurs in the phagosome of activated neutrophils and involves the production of highly antimicrobial RCS/ROS to damage and kill the engulfed pathogen. Our work in clinical isolates revealed a strong correlation between increased HOCl resistance and the presence of *rcrB,* independent of whether the strain was isolated from patients with asymptomatic or symptomatic infections in either the lower (i.e. cystitis) or upper (i.e. pyelonephritis) urinary tract (**FIG 2**). Whether UPEC’s ability to express RcrB in response to HOCl, an oxidant prominent during inflammation, provides a survival benefit *in vivo* is an interesting question that we would like to answer in the future. Deciphering the strategies that bacterial pathogens utilize to defend themselves against the deleterious effects of ROS/RCS is therefore important to better understand their pathogenesis. Moreover, it may provide new perspectives on potential novel drug targets, such as RcrB, which could impair pathogen survival in this highly oxidizing environment and thus increase the efficacy of the immune system. This becomes increasingly pressing in times of increasing antibiotic resistance mechanisms are reported for UPEC. Further, we showed that protection by RcrB is provided in actively growing as well as dormant cells and occurs independent of the growth phase (**FIG 2**). Limitation appears to be that cells must be in their planktonic state, as the presence of RcrB did not affect UPEC’s biofilm forming abilities (**FIG 4**). While UPEC form biofilms, these occur mostly intracellularly after invasion of uroepithelial cells, which presents a more protective environment and shields them from extracellular stress (48). Although the precise biological functions of *rcrA* and *rcrB* are yet to be elucidated, it is now evident that at least RcrB is a major player for bacterial HOCl resistance *in vitro* and that its function likely is conserved across several *E. coli* pathotypes and possibly different bacterial species.

Besides methionine and thiol groups, HOCl shows high reactivity towards amino groups, resulting in the formation of chloramines (4). However, it is likely that protein chloramines decompose into smaller chloramines and/or aldehydes, which are more stable and readily penetrate the bacterial cell. Moreover, chloramines formation is also a result of the reaction between HOCl and ammonium, glycine, and/or the non-proteinogenic amino acid taurine (43–46). Despite having lower reactivities than HOCl, chloramines still show prolonged bactericidal properties and account for the majority of the added HOCl (50). Indeed, it has been shown that tyrosine chlorination was more pronounced in phagosomal proteins compared to proteins of ingested bacteria (51), suggesting that phagosomal proteins are exposed to a higher HOCl concentration and that chloramines may be the more physiological species that pathogens experience. Whether RCS other than HOCl have similar effects on the expression of the RcrR regulon has not yet been studied. In our current study, we show that *rcrARB* upregulation is induced by several members of the RCS family, including Ca(OCl)_2_, NH_2_Cl, Gly-Cl, and NCT (**FIG 7**). Moreover, *rcrARB* expression is also induced by HOBr, an equally powerful yet physiologically less relevant oxidant (52). Our data therefore suggest that each of these RCS can effectively oxidize RcrR and cause the necessary conformational changes in the repressor that result in elevated RcrB expression and UPEC survival (**FIG 7&8**). While we did not observe any significant differences in the extent to which the *rcrARB* operon was induced by the individual RCS, RcrB was most protective during HOCl stress, while its effect was less pronounced in the presence of NH_2_Cl, Gly-Cl, and NCT (**FIG 7&8**). We observed upregulation of *rcrARB* and protection by RcrB in the presence of NCT, a product of the chlorination of taurine after HOCl exposure, which is unable to penetrate through the plasma membrane. However, we have performed our studies in MOPS glucose minimal media, which contains ammonia in sufficiently high concentrations (53) to allow the conversion of NCT into NH_2_Cl (43), which readily enters the cytoplasm. However, neither did we observe any significant upregulation of *rcrARB* nor any impact of RcrB on survival when UPEC was exposed to H_2_O_2_ and HOSCN (**FIG 7&8**). Both oxidants are characterized by their high thiol-specificity, which makes them distinctly different from RCS (54). A study in *Pseudomonas aeruginosa* revealed overlapping outcomes for treatments with HOCl and HOBr as both oxidants target non-growing cells more efficiently than growing cells and elicit similar bacterial responses (52). In contrast, treatment with HOSCN has been found to affect primarily actively growing cells and evoking different defense mechanisms, likely due to its highly thiol-specific nature (52). Recent progress in the field of bacterial oxidative stress defense systems was made by the discovery of HOSCN reductases, which efficiently reduce HOSCN. The presence of these enzymatic systems potentially provide an explanation why RcrR does not sense thiol-specific oxidants such as HOSCN or H_2_O_2_ (24, 55–58).

Proteins constitute for >50% of the cellular macromolecules and are known to rapidly react with HOCl (59). Numerous studies in different HOCl-treated bacterial species revealed the strong upregulation of the heat shock regulon, indicating an accumulation of misfolded proteins and supporting the idea that proteins are the major targets of HOCl (28, 52, 60–63). Similarly, H_2_O_2_ can cause substantial protein aggregation as the result of methionine and cysteine oxidation (64). It is therefore not surprising that bacteria have evolved a number molecular chaperones as defense mechanisms to limit and repair RCS-mediated damage, such as Hsp33 (65), CnoX (66, 67), RidA (68, 69), and polyP (70). Intriguingly, inhibitors of polyP synthesis have been shown to be effective against UPEC and reduce the bacterial load during acute UTI (71, 72). Moreover, microbial responses to ROS/RCS often involve redox-sensitive transcriptional regulators, which use conserved cysteine and/or methionine residues to modulate their activity (73). This, in turn, upregulates the transcription of their target genes, many of which have been shown to protect the organism from ROS/RCS. Three RCS-responsive transcriptional regulators have been identified in *E. coli*, all of them in the K12-strain MG1655: *(i)* HypT (26); *(ii)* the TetR-family transcriptional repressor NemR (60) and *(iii)* the AraC-family transcriptional activator RclR (74, 75). NemR is a transcriptional repressor that is inactivated by HOCl, resulting in the expression of *nemA*, an N-ethylmaleimide reductase, and *gloA*, a glyoxylase, both of which have been shown to contribute to HOCl resistance in *E. coli* (21). The transcriptional activator RclR is specifically sensitive to RCS and HOSCN and regulates the *rclABC* gene cluster (24, 25, 75). While the function of RclB and RclC are still unknown, RclA was identified as a Cu(II) reductase that protects *E. coli* from HOCl stress and, more recently, an HOSCN reductase (24). With the RcrR regulon, we have added another regulatory system to the list of RCS-sensing transcriptional regulators (28). Questions remained as to why UPEC requires the presence of three regulons (i.e. RcrR, NemR, and RclR) to defend against HOCl stress. We found that all three systems were upregulated after just 5 minutes of exposure to HOCl (**FIG 6**). While *nemR/nemA* and *rclR/rclA* expression started to decline after 10 minutes, *rcrARB* expression remained unaffected throughout the 45 minutes tested. Further, our LPE assays indicate that Δ*rcrB* mutants are much more susceptible to HOCl than cells that are defective in a member of the two other regulons (**FIG 9**) (28). Likewise, RcrB expression significantly increased the HOCl resistance of a Δ*rclC* mutant, while the expression of RclC had no additional protective effect on Δ*rcrB*, suggesting that both membrane proteins are functionally different despite their sequence homology. While expression of *rclABC* was induced following phagocytosis by neutrophils, the Δ*rclABC* mutant did not show any differences in neutrophil killing compared to the wildtype (42), which is in stark contrast to a Δ*rcrB* mutant, which was significantly more sensitive to neutrophil killing compared to the corresponding wildtype CFT073 (28). This shows the substantial role of RcrB in resistance to both HOCl and neutrophils compared to the other stress defense systems. The bacterial response and defense strategies are expected to be critical for their ability to survive the immune cell attack, as reported in a recent review of several studies (17). Moreover, independent studies confirmed that the presence of functional oxidative stress defense systems positively affects pathogen colonization in the host, emphasizing their importance for pathogenesis (71, 72, 76, 77).

## MATERIAL AND METHODS

### Strains, plasmids, oligonucleotides, and growth conditions

All strains, plasmids, and oligonucleotides used in this study are listed in Table 1. Unless otherwise mentioned, bacteria were cultivated at 37 °C and 300 rpm in luria broth (LB, Millipore Sigma) or in 3-(N-morpholino)propanesulfonic acid minimal media containing 0.2% glucose, 1.32 mM K_2_HPO_4_ and 10 μM thiamine (MOPSg) (53). Kanamycin (100 μg/ml), ampicillin (200 μg/ml), or chloramphenicol (34 μg/ml) were added when required.

### Construction of in frame gene deletions in CFT073 and *E. coli* Nissle 1917

In-frame deletion mutants were constructed using the lamda red-mediated site-specific recombination (78). CFT073 genes *rclC, rcrR*, and *rcrB* as well as the *rcrB* homologous gene in *E. coli* Nissle 1917 were replaced with a chloramphenicol resistance (Cm^R^) resistance cassette, which was resolved using pCB20 to yield the nonpolar in-frame deletion strains CFT073Δ*rcrB*Δ*rclC,* CFT073Δ*rclC*, CFT073Δ*rcrR*, CFT073Δ*rcrB*, and EcNΔ*rcrB-H,* respectively (78). All chromosomal mutations were confirmed by PCR.

### Plasmid construction

The *rcrB* and *rclC* gene were amplified from UPEC strain CFT073 genomic DNA with primers listed in Table S1 and cloned into the *Nde*I and *BamH*I sites of plasmid pET28a to generate the N-terminally His_6_-tagged RcrB and RclC expression plasmid pJUD29 and pJUD46, respectively. All constructs were verified by DNA sequencing (Eurofins).

### Preparation of oxidants

Sodium hypochlorite (Millipore-Sigma) stock solution was utilized in the preparation of many oxidants. The molar concentration of the stock solution was determined by measuring the *A*_292 nm_ after diluting in 10 mM NaOH and using ε_292_ = 350 M^-1^ cm^-1^. As previously described (42), chloramines were prepared freshly by mixing HOCl with excess ammonium sulfate, glycine, and taurine, respectively. The concentration of chloramines was determined by measuring absorbance after diluting in 10 mM NaOH, using ε_250 nm_ = 429 M^-1^ cm^-1^ (79). The molar concentration of hydrogen peroxide was determined by measuring the *A*_240 nm_ after diluting the stock solution in 50 mM KPi buffer and using ε_240_ = 43.6 M^-1^ cm^-1^. HOSCN was prepared in an enzyme-catalyzed reaction using lactoperoxidase as previously described in (52, 80).

### Studies in artificial urine media (AUM)

AUM was prepared as described in (41) with the following components in their final concentrations: 78.5 mM sodium chloride, 9 mM sodium sulfate, 2.2 mM sodium citrate dihydrate, 0.1 mM sodium oxalate, 3.6 mM potassium dihydrogen phosphate, 21.5 mM potassium chloride, 3 g/L tryptic soy broth, 3 mM calcium chloride, 2 mM magnesium chloride, 15 mM ammonium chloride, 6 mM creatinine, and 200 mM urea (pH 6.3). AUM was prepared fresh every 7-10 days.

### Determining oxidant susceptibility through lag phase extension (LPE) analyses

Overnight cultures of the indicated strains were diluted 25-fold into the indicated media (i.e. MOPSg or AUM) and cultivated until late-exponential / early stationary phase (OD_600_ = ∼2). Cultures were back diluted into fresh MOPSg/AUM to an OD_600_= 0.03 and cultivated in a Tecan Infinite 200 plate reader in the presence or absence of increasing concentrations of the indicated oxidant. To test the role of RcrB in exponentially growing cultures (**FIG 3A, Supplementary FIG S2B**), overnight cultures were diluted into fresh MOPSg to an OD_600_ = 0.01 and grown to the indicated OD_600_ before they were split up and cultivated in the presence of the indicated HOCl concentrations. To test the role of RcrB in late log cultures (**FIG 3B, Supplementary FIG S2A**), overnight cultures were diluted 25-fold into fresh MOPSg and grown until they reached late log / early stationary phase (OD_600_ ∼2) before they were diluted again to the indicated OD_600_ and cultivated in the presence of the indicated HOCl concentrations. A_600nm_ measurements were recorded every 10 min for 16 h. Oxidant sensitivities of the strains tested were examined by quantifying their oxidant mediated LPE. LPE were calculated by determining the differences in time for oxidant-treated samples to reach *A*_600 nm_ > 0.3 compared to the untreated controls as described in (25, 28).

### Gene expression analyses using quantitative real-time PCR (qRT-PCR)

Overnight LB cultures of the indicated strains were diluted into MOPSg to an *A*_600 nm_= 0.1 and cultivated until OD_600_= 0.5-0.55 was reached before they were either left untreated or treated with the indicated concentrations of oxidant. At the indicated time points, 1 ml cells were harvested onto 1 ml of ice-cold methanol to stop transcription. After centrifugation, total RNA was prepared from the cell pellet of three biological replicates of the untreated and oxidant-treated strains using a commercially available RNA extraction kit (Macherey & Nagel). Remaining DNA was removed using the TURBO DNA-free kit (Thermo-Scientific) and cDNA generated using the PrimeScript cDNA synthesis kit (Takara). qRT-PCR reactions were set up according to the manufacturer’s instructions (Alkali Scientific). Transcript level of the indicated genes were normalized against transcript level of the 16S rRNA-encoding *rrsD* gene and relative fold-changes in gene expression were calculated using the 2^-ΔΔC^_T_ method (81).

### Bacterial survival assay after exposure to HOCl

Unless stated otherwise, pre-cultures of the UPEC strains CFT073 and Δ*rcrB* were grown aerobically at 37°C for 16-18 h in LB. For the survival analyses of actively growing cells, the overnight cultures were diluted into MOPSg media to an OD_600_= 0.01 and cultivated until OD_600nm_ reached 0.1. 1 mL of cells were pelleted at 3,000 x *g* for 5 min, washed twice and then resuspended in pre-warmed MOPS (no glucose, non-growing) and MOPSg (actively growing). 190 µL of the resuspended cultures were immediately transferred into a pre-warmed 96-well plate containing the indicated concentrations of HOCl and incubated aerobically at 37 °C. For stationary cultures, the indicated strains were grown in MOPSg media for ∼22 h before they were pelleted and resuspended in MOPS (no glucose, non-growing) and MOPSg (actively growing) to an OD_600nm_= 0.1. Cultures were then incubated in 96-well plates in the presence/ absence of HOCl-stress for 145 min, serially diluted in PBS, and spotted onto LB agar plates. CFU counts were examined after 16 h of incubation.

### Biofilm formation in presence of HOCl

The effect of HOCl on UPEC biofilm formation was examined for CFT073 and Δ*rcrB* following a previously published protocol (39) with minor modifications. Overnight LB cultures were diluted into LB without salt to an OD_600_= 0.05. 150 µL of cultures then transferred into polystyrene 96-well plates and incubated statically at 37°C to allow for attachment. After six hours, HOCl was added at the indicated concentrations, the plate was gently rotated to mix HOCl into the culture and incubated for additional 16 h under the same static conditions. Next, OD_600_ was measured, and the planktonic cells were removed. The biofilms were washed twice with PBS, stained with 0.1% crystal violet (CV) for 15 min, washed again twice with PBS to remove unbound CV dye. CV bound to biofilm cells was extracted with 30% (v/v) glacial acetic acid and OD_595_ was determined.

### Time Killing Assays

Overnight MG1655, CFT073, and Δ*rcrB* cultures were diluted ∼25-fold into AUM by normalizing cultures to an OD_600_= 0.1 in a 6-well sterile cell culture plate. HOCl was added at a final concentration of 664 μM and the plates incubated at 37°C at 150 rpm under aerobic conditions. After 60 min, OD_600_ of cultures were recorded, sample volumes were normalized to the lowest optical density measured, serially diluted in PBS (pH = 7.4) and plated on LB agar to quantify colony forming units (CFU) after 16 hours. Percent survival was calculated as the ratio of surviving colonies from treated to untreated samples.

### Oxidant susceptibility determined by an overnight growth assay and end point OD_600_ measurement

The assay was performed as described for the LPE assay with the only exception that the 96-well plate was incubated in a shaking incubator at 37°C and 300 rpm. One final OD_600_ measurement was recorded after 16-18 h.

## ACKNOWLEDGEMENTS

This work was supported by the NIAID grant R15AI164585 and the Illinois State University Pre-Tenure Faculty Initiative Grant (to J.-U. D.). M.E.C was supported by a Weigel fellowship by the Phi-Sigma Biological Sciences Honors Society and the Jill Hutchison endowed scholarship. L.F.G. and L.P. were supported by a DAAD RISE fellowship from the German Academic Exchange Service. M.B. and B.R. received FireBIRD grants by the Illinois State University Undergraduate Research Support Program. We thank Dr. Maria Hadjifrangiskou (Vanderbilt University) for providing the UPEC clinical isolates as gifts and Dr. Michael J. Gray (University of Alabama) for the *E. coli* Nissle 1917 strain. Members of the Dahl lab are acknowledged for feedback and proof-reading of the manuscript.

## Supplemental Figures

**Supplementary FIG S1.**
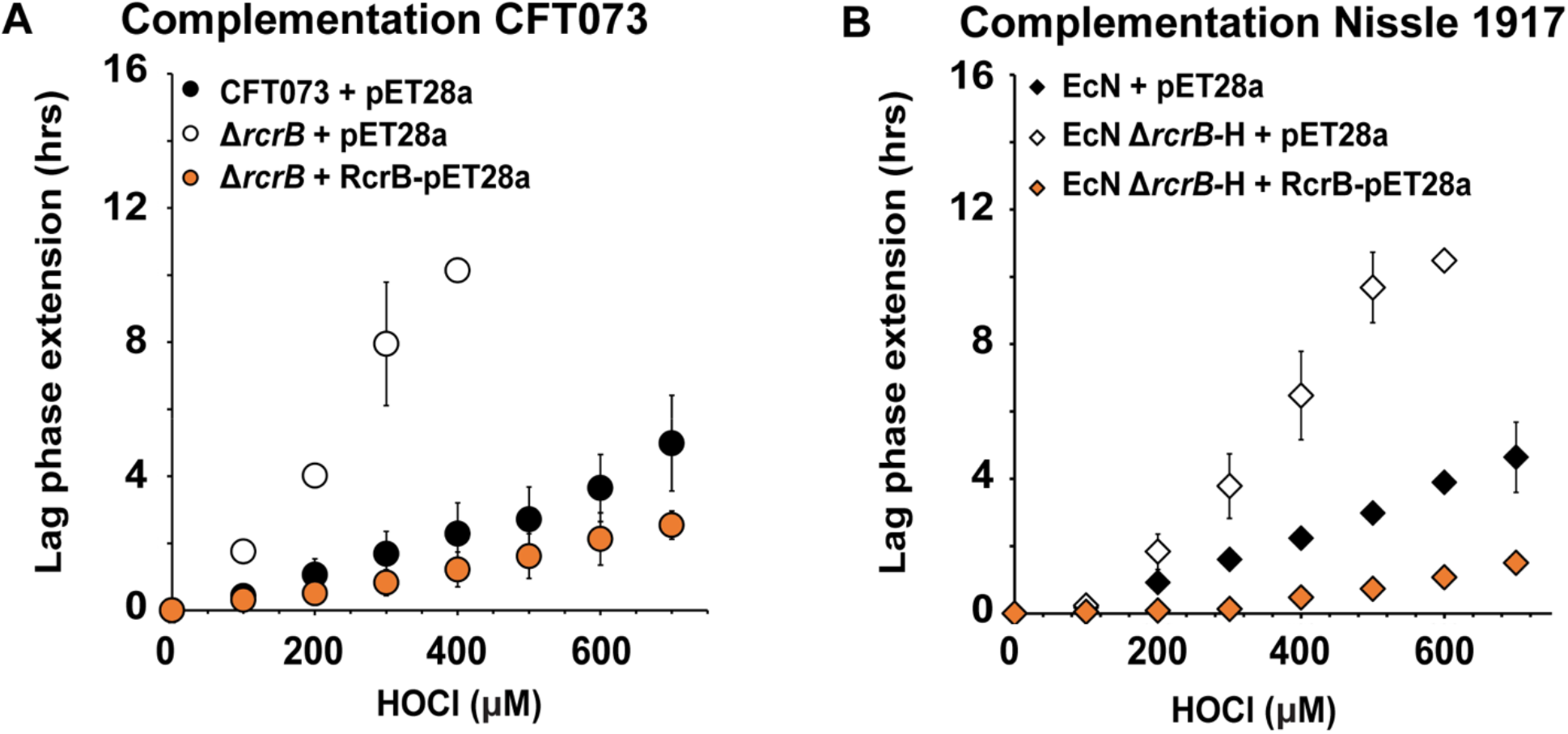
Recombinant expression of CFT073 RcrB complements the HOCl sensitivity phenotypes of *E. coli* strains deficient in *rcrB* or its homolog. Growth phenotype analyses of the UPEC strain CFT073 and Nissle 1917 carrying pET28a or *rcrB*-pET28a in the presence and absence of *rcrB* were performed in MOPSg media in the presence of the indicated HOCl concentrations. HOCl-mediated LPE was calculated for each strain (see Materials and Methods for a detailed protocol). Recombinant expression of RcrB in (A) CFT073 and Δ*rcrB*, and (B) *E. coli* Nissle 1917 and Δ*rcrB*-homolog (Δ*rcrB*-H) resulted in full complementation restoring HOCl resistance comparable to the corresponding wildtype strains CFT073 and EcN, respectively *(n = 4, ± S.D.)*.

**Supplementary FIG S2.**
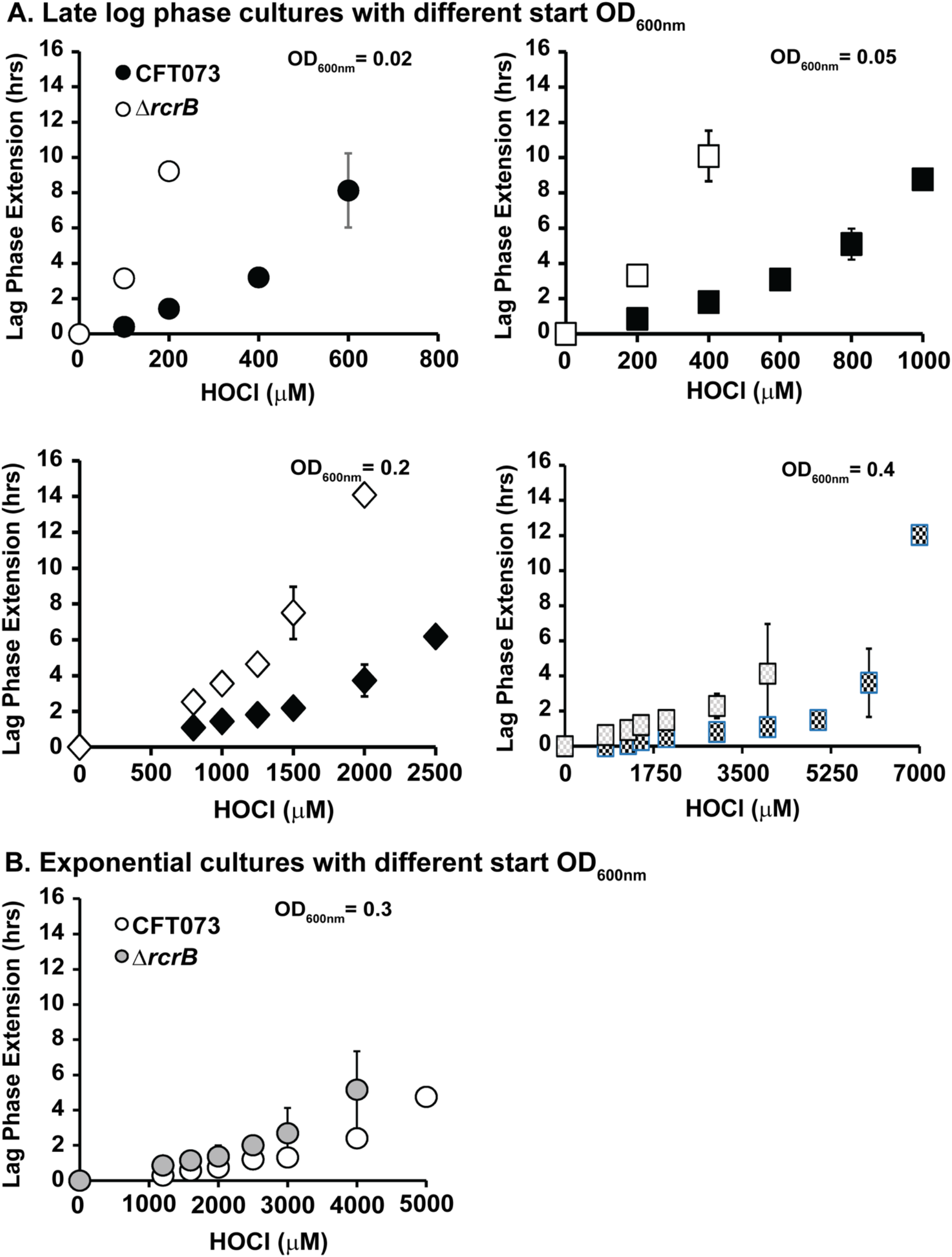
RcrB plays an important role for UPEC’s HOCl resistance under various growth conditions. Growth phenotype analyses of UPEC strains CFT073 and Δ*rcrB* were performed in MOPSg media in the presence of the indicated HOCl concentrations. HOCl-mediated LPE was calculated for each strain (see Materials and Methods for a detailed protocol). **(A)** *Late log cultures:* overnight cultures of CFT073 (black fillings) and Δ*rcrB* (white fillings) were diluted 25-fold into fresh MOPSg and grown until they reached late log / early stationary phase (OD_600_ ∼2) before they were diluted again to the indicated start OD_600_ and cultivated in the presence of the indicated HOCl concentrations; *(n = 3-4, ± S.D.)*. **(B)** *Exponential cultures:* overnight cultures of CFT073 (white circles) and Δ*rcrB* (grey circles) were diluted into fresh MOPSg to an OD_600_ = 0.01 and grown until they reached an OD_600_ = 0.3 before they were split up and cultivated in the presence of the indicated HOCl concentrations; *(n = 3-4, ± S.D.)*.

**Supplementary FIG S3.**
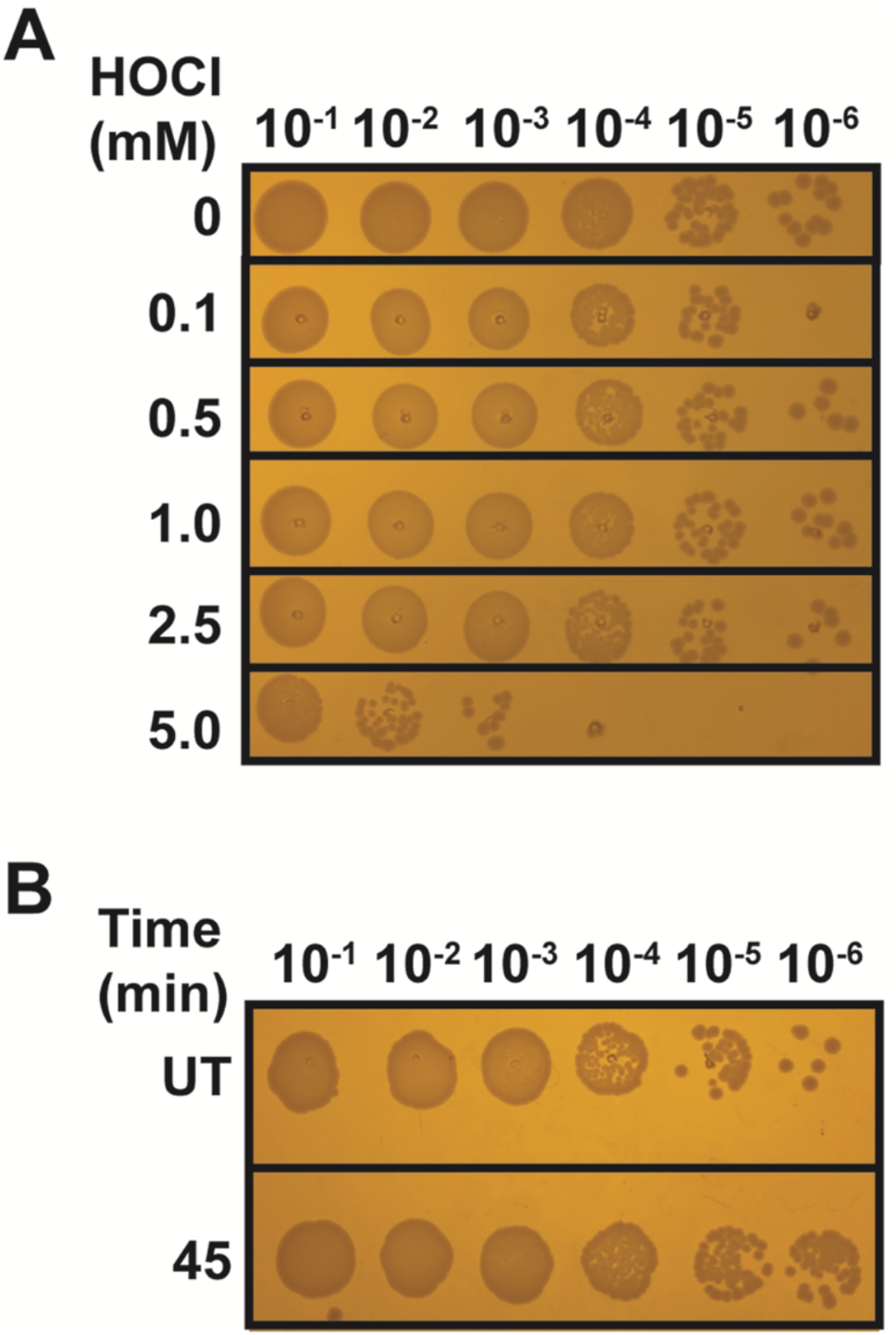
Survival assays performed in conjunction with gene expression analyses. UPEC strain CFT073 was grown in MOPSg to mid-log phase (OD_600_ ∼0.5-0.55) before it was incubated with the indicated HOCl concentrations for the indicated time. Remaining HOCl was quenched with 5-fold excess of thiosulfate after **(A)** 15 min and **(B)** 45 min of exposure to 1 mM HOCl, cells serially diluted and spot-titered onto LB agar plates for overnight incubation at 37 °C. The experiments were repeated at least three independent times.

**Supplementary FIG S4.**
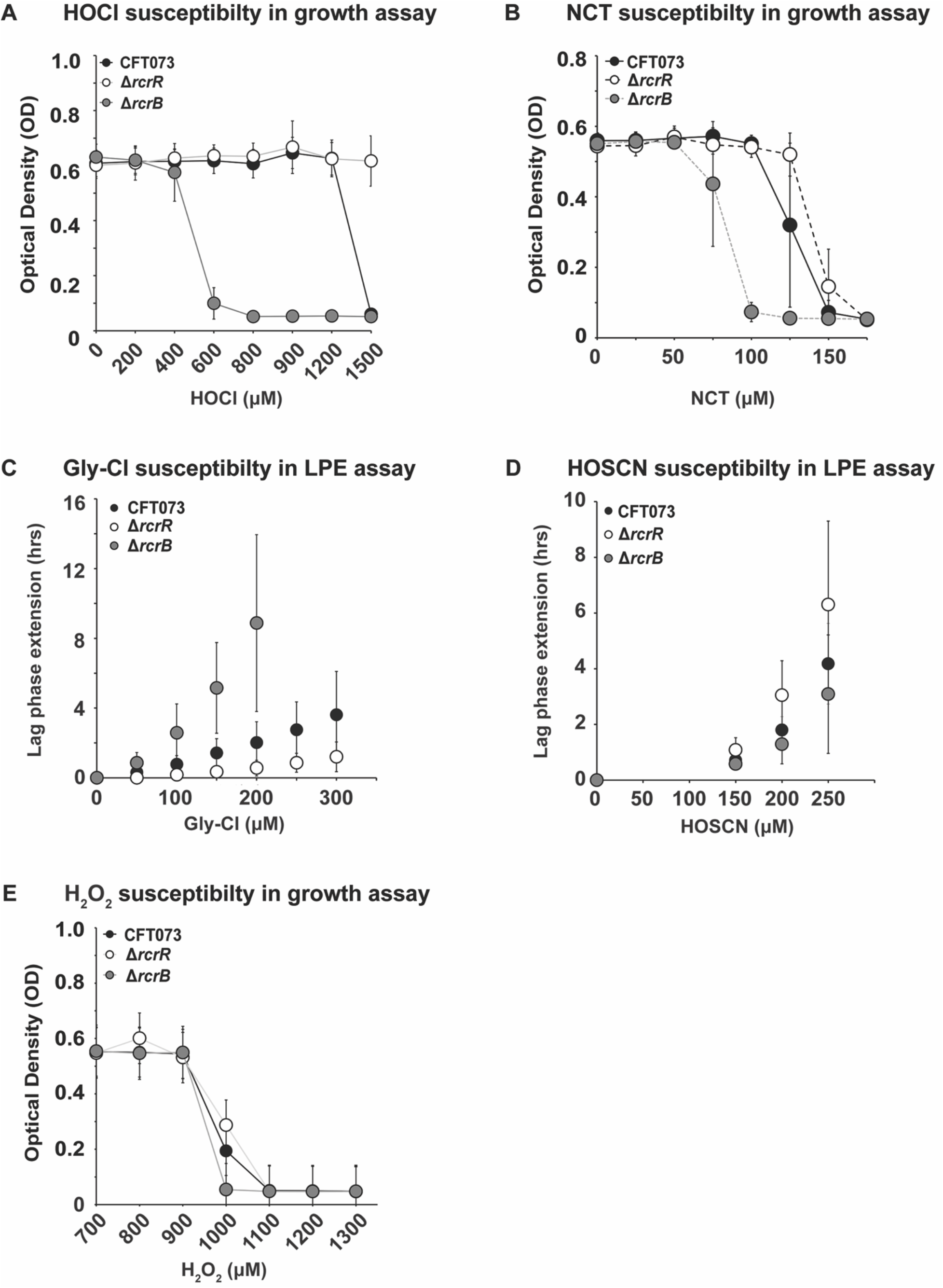
The RcrR regulon protects from HOCl stress and its byproducts but not from H_2_O_2_ and HOSCN. Growth phenotype analyses of UPEC strains CFT073, Δ*rcrB*, and Δ*rcrR* were performed in MOPSg media in the presence of the indicated stressor concentrations. LPE were calculated for each strain (see Materials and Methods for a detailed protocol). The absence of *rcrB* in CFT073 caused a reduction in HOCl resistance, while constitutive RcrB expression (i.e. Δ*rcrR*) caused increased resistance to all stressors except HOSCN and H_2_O_2_, *(n = 4-6, ± S.D.)*.

**Supplementary FIG S5.**
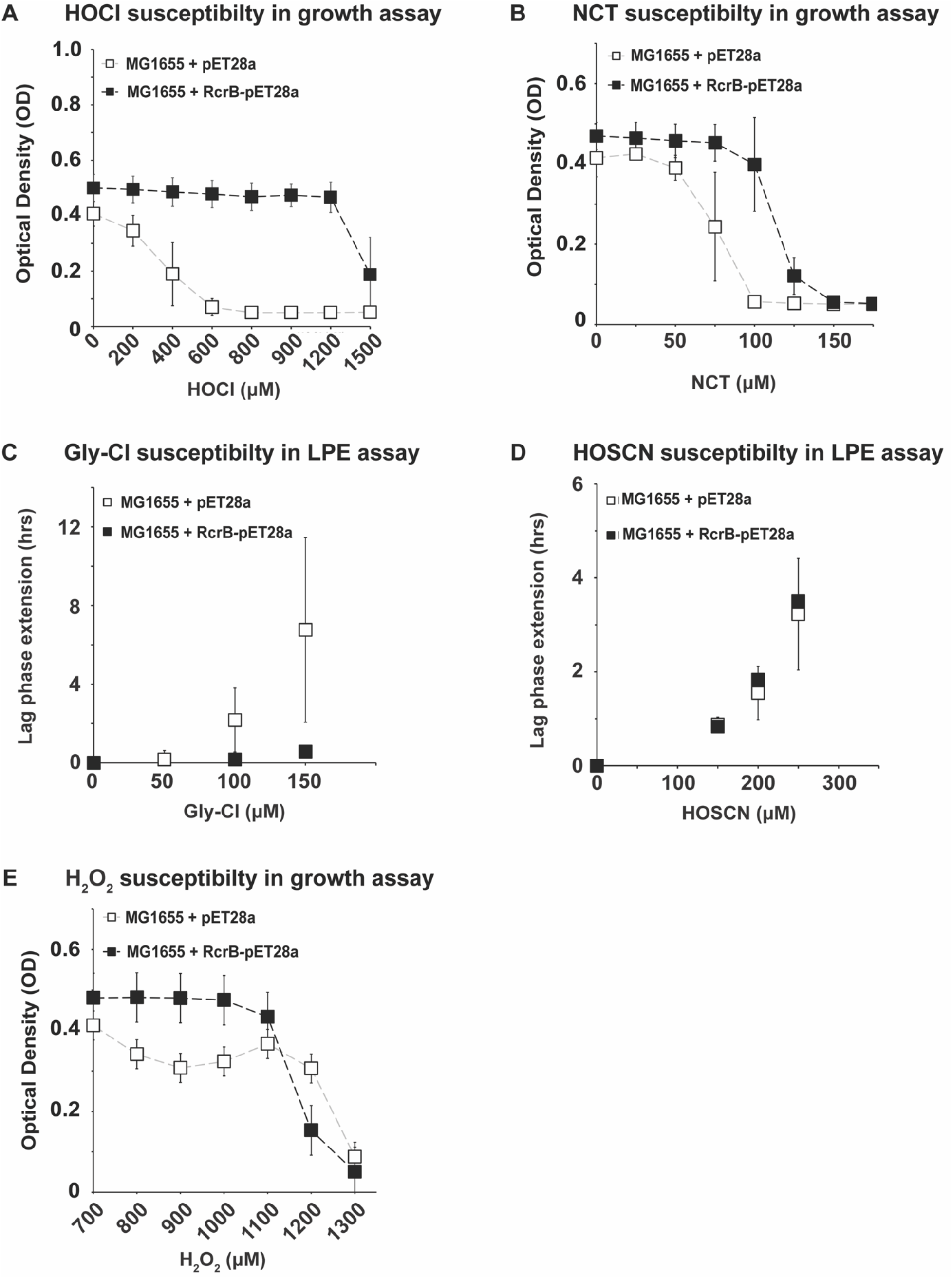
The RcrR regulon protects from HOCl stress and its byproducts but not from H_2_O_2_ and HOSCN. Growth phenotype analyses of the K-12 *E. coli* strain MG1655 carrying pET28a or *rcrB*-pET28a, respectively, were performed in MOPSg media in the presence of the indicated stressor concentrations. LPE were calculated for each strain (see Materials and Methods for a detailed protocol). Recombinant expression of RcrB in MG1655 resulted in an increased resistance to all stressors except for HOSCN and H_2_O_2_ compared to the empty vector control, *(n = 4-6, ± S.D.)*.

